# The plasma Factor XIII heterotetrameric complex structure: unexpected unequal pairing within a symmetric complex

**DOI:** 10.1101/651448

**Authors:** Sneha Singh, Alexis Nazabal, Senthilvelrajan Kaniyappan, Jean-Luc Pellequer, Alisa S. Wolberg, Diana Imhof, Johannes Oldenburg, Arijit Biswas

## Abstract

Factor XIII (FXIII) is a predominant determinant of clot stability, strength, and composition. Plasma FXIII circulates as a pro-transglutaminase with 2 catalytic A subunits and 2 carrier-protective B subunits in a heterotetramer (FXIII-A_2_B_2_). FXIII-A_2_ and -B_2_ subunits are synthesized separately and then assembled in plasma. Following proteolytic activation by thrombin and calcium-mediated dissociation of the B-subunits, activated FXIII (FXIIIa) covalently cross-links fibrin, promoting clot stability. The zymogen and active states of the FXIII-A subunits have been structurally characterized; however, the structure of FXIII-B subunits and the FXIII-A_2_B_2_ complex have remained elusive. Using integrative hybrid approaches including atomic force microscopy, cross-linking mass spectrometry, and computational approaches, we have constructed the first all-atom model of the FXIII-A_2_B_2_ complex. We also used molecular dynamic simulations in combination with isothermal titration calorimetry to characterize FXIII-A_2_B_2_ assembly, activation, and dissociation. Our data reveal unequal pairing of individual subunit monomers in an otherwise symmetric complex, and suggest this unusual structure is critical for both assembly and activation of this complex. Our findings enhance understanding of mechanisms associating FXIII-A_2_B_2_ mutations with disease and have important implications for the rational design of molecules to alter FXIII assembly and/or activity to reduce bleeding and thrombotic complications.

## INTRODUCTION

Plasma coagulation factor XIII (FXIII) circulates as a heterotetramer composed of two catalytic FXIII-A subunits tightly-associated (10^−7^ - 10−^9^ M) with two carrier/regulatory FXIII-B subunits (FXIII-A_2_B_2_)(1, 2). During coagulation, proteolytic activation by thrombin and calcium-mediated dissociation of FXIII-B subunit generates activated FXIII-A (FXIIIa) that covalently cross-links fibrin, promoting clot stability(3). Deficiency in plasma FXIII antigen or activity is associated with mild-to-severe bleeding(3).

The structure of the catalytic FXIII-A subunits is well-characterized, consisting of an activation peptide (FXIII-AP; residues 1-37) followed by four distinct domains: β-sandwich (residues 38-183), central core (residues 184-515), β-barrel-1 (residues 516-627), and β-barrel-2 (residues 628-731)(4). Both zymogen and activated forms of FXIII-A have been crystallized(4, 5), revealing a compact structure of zymogen FXIII-A_2_, but an open, extended conformation of activated FXIIIa. Despite the essential regulatory role of FXIII-B, structural information on this molecule is sparse. Sequence homology with complement proteins suggest each FXIII-B subunit is composed of ten sushi domains, each containing ~60 amino acid residues and two disulfide bonds(6). Although sedimentation analysis initially suggested FXIII-B is a monomer, more recent data suggest FXIII-B circulates as a dimer(6, 7). Sushi domains 4 and 9 (S4 and S9) are thought to mediate FXIII-B_2_ dimerization, whereas S1 and S2 are thought to promote interactions with the FXIII-A subunits(1, 6). The size and complexity of FXIII-A_2_B_2_ make it difficult to characterize by traditional methods such as X-ray crystallography or NMR. Apart from a partial all-atom model generated with minimal experimental data(8), there is no detailed model for the FXIII-B_2_ dimer or the FXIII-A_2_B_2_ complex. Consequently, knowledge of the structural interface between FXIII-B subunits or the FXIII-A_2_B_2_ heterotetramer is incomplete. Structural resolution of plasma FXIII-A_2_B_2_ and its transition to activated FXIIIa is essential for defining implications of missense FXIII mutations, as well as the development of potential FXIII(a) inhibitors for treating bleeding and thrombotic complications associated with abnormal clot stabilization

Integrative/hybrid (IH) approaches are useful for dissecting the structural architecture of complexes that escape traditional structural determination techniques(9, 10). These approaches integrate biochemical and computed data to yield structural information on macromolecular complexes. For example, IH has revealed detailed molecular conformational states of glucagon receptor(11) and the chromatin remodeling complex(12), and recently a combination of structural methods with AFM has provided key information on factor Va bound to activated protein C(13). Since FXIII-A_2_B_2_ has not been amenable to traditional structural analysis, we addressed this gap using a bootstrapped IH approach. We first used atomic force microscopy to define the macromolecular structure of the FXIII subunits individually and as a complex, and chemical crosslinking/mass spectrometry (XL-MS) to define residues in the FXIII-A_2_B_2_ intersubunit interface. We then used these data as structural constraints to assemble first a monomeric FXIII-B subunit model followed by an all-atoms structural model of the FXIII-A_2_B_2_ complex. We then overlaid these putative models on surface topographic atomic force micrographs of FXIII-A_2_B_2_ to produce a complete macromolecular structure^10^. Finally, we integrated molecular simulations from the all-atoms model with isothermal calorimetry to interrogate conformational dynamics during FXIII-A_2_B_2_ assembly, activation, and subunit disassociation. Our data indicate FXIII-A_2_B_2_ assembles with unexpected unequal pairing within an otherwise symmetric complex and suggest this conformation is essential for FXIII function. These findings provide the first molecular structure of this important coagulation protein.

## MATERIAL AND METHODS

Software, databases, and webservers used in this study are listed in *Supplemental Table 1*. A flowchart for the IH approach is presented in *Supplemental Figure 1*. A conformational ensemble of structures from the post-equilibration simulation trajectory of heterotetramer complex model 1 is submitted to the PDB-dev IH structure database(14, 15). Detailed methodology can be found in the *Supplemental methods*.

### FXIII-A and FXIII-B subunit, cloning expression and purification

Cloning expression and purification of rFXIII-A (recombinant FXIII-A) subunit was performed as described by Gupta et al, 2016 (8)(Supplemental Figure 2a). Human FXIII-B cDNA, inserted into the cloning site of pReciever-M01 mammalian expression vector was transfected into *HEK293t* cells as per previously reported protocol(16). Secreted protein harvested posttransfection was concentrated and subjected to immunoaffinity purification using the Thermo Scientific Pierce Co-IP kit (Pierce Biotechnology, USA) (Supplemental Figure 2b).

### Purification of FXIII-A_2_B_2_ complex

FibrogamminP (CSL Behring, Germany) reconstituted in water was run on a Superdex 200 increase 10/300 column /Äkta Pure system (GE healthcare, Germany) equilibrated with 20mM Tris, 100mM NaCl at pH 7.4. Peak corresponding to FXIII-A_2_B_2_ (320Kda), was re-purified thrice until a highly pure, single, homogenous, monodispersed peak was obtained with no excipients (Supplemental Figure 2c). The eluted peak was concentrated and quantified for further downstream applications.

### Atomic force microscopy (AFM) of FXIII

Surface topology of recombinant FXIII-A_2_ (rFXIII-A_2_ expressed and purified in-house), recombinant FXIII-B_2_ (rFXIII-B_2_ expressed and purified in-house), and FXIII-A_2_B_2_ [purified from FibrogamminP (CSL Behring, Marburg, Germany)] was analyzed individually using high-resolution AFM. Briefly, samples suspended in PBS pH 7.4 (1–2 mM) were placed on a freshly-cleaved mica and allowed to adsorb (15 minutes). Non-adherent proteins were removed by washing twice with imaging buffer (10 mM Tris-HCl; 50 mM KCl, pH 7.4). Samples were imaged in oscillation mode in liquid (imaging buffer) as described(17) and acquired using the Nanoscope III microscope.

### XL-MS of FXIII-A_2_B_2_ heterotetramer complex

One μL of 3.12 μg/mL purified FXIII-A_2_B_2_ was mixed with 1 μL of a matrix of re-crystallized sinapinic acid (10 mg/mL) in acetonitrile/water (1:1, v/v), triflouroacetic acid (TFA) 0.1% (K200 MALDI Kit; CovalX, Zurich, Switzerland). After mixing, 1 μL of each sample was spotted on the MALDI plate. After crystallization at room temperature, the plate was introduced in the MALDI mass spectrometer (Ultraflex III MALDI ToF, Bruker Daltonik GmbH, Bremen, Germany) equipped with HM2 High-Mass detection (CovalX, Zurich, Switzerland) and analyzed immediately in High-mass MALDI mode. MS data were analyzed using Complex Tracker analysis software (CovalX, Zurich, Switzerland). For characterization and peptide mass fingerprinting, the purified FXIII-A_2_B_2_ complex was subjected to ASP-N, trypsin, chymotrypsin, elastase, and thermolysin proteolysis, followed by nLC-LTQ Orbitrap MS/MS analysis (formic acid 1% added to the final solution after digestion). Purified FXIII-A_2_B_2_ (1.25 μM) was crosslinked with 2 μL of DSS (d0d12) reagent (Creative Molecules Inc., Canada) at room temperature for 3 hours, prior to digestion. Nano-LC chromatography was performed using an Ultimate 3000 (Dionex, IL, USA) system in-line with an LTQ Orbitrap XL mass spectrometer (ThermoFischer Scientific, IL, USA). Acquired data were analysed by XQuest version 2.0 and Stavrox version 2.1. The FXIII-B intra-subunit and FXIII-A-FXIII-B inter-subunit cross-linked peptides and residues are presented in *Supplemental Tables 2* and *3*.

### Generation of the FXIII-B subunit model

FXIII-B intra-subunit XL-MS crosslinked residues were matched to residue contact prediction data to generate constrained models of FXIII-B monomers on the AIDA server (http://aida.godziklab.org/)(18) (*Supplemental Figures 3 and 4*). Sushi domains were based on previously-generated high-quality threaded models from I-TASSER(19) (https://zhanglab.ccmb.med.umich.edu/I-TASSER/ (*Supplemental Figures 5a-5j*). We also assembled a FXIII-B subunit monomer model (*Supplemental Figure 4*) in default mode, i.e. without constraints and docked this model symmetrically (M-Z docking server(20)) to model unbound FXIII-B_2_ dimer.

### Generation of the FXIII-A_2_B_2_ all-atom model

Inter-subunit, XL-MS-directed docking of all FXIII-B monomer conformations on the FXIII-A_2_ crystal structure (PDB ID: 1f13) was performed using the HADDOCK expert interface webserver (http://milou.science.uu.nl/services/HADDOCK2.2/)(21). Since this webserver allows for only bi-molecular docking, whereas the *in-silico* model involves 3 proteins (FXIII-B monomer and FXIII-A_2_ dimer), we treated the dimer as a single molecule by renumbering the residues of each FXIII-A monomer in continuum. We based structural constraints for modeling and docking FXIII-B monomer on FXIII-A_2_ on inter- and intra-subunit cross-linked residues (*Supplemental Tables 2 and 3*). Docking constraints (n=64) required that all residues belong to detected cross-linked peptides that can form side chain contacts (*Supplemental Table 4*) to cover the FXIII-A_2_ /FXIII-B trimer surface. Moreover, FXIII-A_2_/FXIII-B contact residues were assigned constant lower and upper limit distances of 3 and 24 Å, respectively(22). We then manually constructed the resulting docked trimer into a tetramer with bilateral symmetry.

### Molecular dynamic simulations of the FXIII-A_2_B_2_ heterotetramer models

Stability of the top-scoring FXIII-A_2_B_2_ complex (best HADDOCK scores amongst the major docking clusters; *Supplemental Figure 6*) from the HADDOCK(23) server was assessed using all-atoms molecular dynamics (MD) simulations (YASARA Structure suite 17.4.17 platform(21, 23, 24) with the embedded md_sim macro)(25, 26). A steered molecular dynamics (SMD) simulation was separately performed on MD-equilibrated model 1 to dissociate the FXIII-B_2_ subunit dimer from the FXIII-A_2_ dimer. The SMD was performed with md_runsteered macro embedded in YASARA, with minor modifications in the steering force (applied acceleration: 100□pm/ps^2^). Analyses of simulation variables, model quality, and model characteristics are detailed in *Supplemental material*. All subsequent structural analyses were performed on the MD-equilibrated complex model 1.

### Modelling transition states between the first FXIII-A_2_: FXIII-B_2_ contacts and the final FXIII-A_2_B_2_ complex

To generate a model of the initial contact between dimeric FXIII-A_2_ and FXIII-B_2_, we docked the crystal structure of FXIII-A_2_ dimer with the dimeric model of unbound FXIII-B_2_ on the Z-dock rigid docking server(27). We considered the highest scoring complex as the initial contact structure and the FXIII-A_2_B_2_ complex model 1 generated on HADDOCK(21, 23, 24) as the final structure, and submitted these to the MINACTION path server (http://lorentz.dynstr.pasteur.fr/suny/submit.php?id0=minactionpath#submit) to generate C_α_-backbone models of the transition-states between these two structures(28). Once a large number of coarse-grained intermediates were generated, we converted 8 intermediates to full atom models, as described^4^.

### Fitting and docking atomic protein structures on AFM surface topographs

We docked three-dimensional heterotetramer coordinates within the AFM-derived topographic surface (envelope) using the AFM-Assembly protocol(29, 30). We defined the docking score as the number of atoms from the protein structure in the favorable layer, and translated this score into pseudo-energy values, where the best score corresponds to the lowest energy. We ran docking protocols on HADDOCK top-scored FXIII complex models, as well as on the crystal structure of FXIII-A_2_ and the models of unbound FXIII-B_2_. For each docking simulation, we retained the top 10^5^ potential solutions and further analyzed the top 10 to produce the minimum docking energy, average energy of the top 10 docking solutions, root-mean-square deviation (RMSD) of the top 3 docking solutions and shift of the best docking solution from the center of the docking grid.

### ITC-based thermodynamic profiling of the assembly and dissociation of the FXIII-A_2_B_2_ heterotetramer complex

Finally, we directly measured thermodynamic changes during complex assembly and disassembly of the FXIII-A_2_B_2_ heterotetramer using ITC on a MicroCal200 microcalorimeter (Malvern Panalyticals, Malvern, UK). To examine FXIII subunit association, we titrated 2.5 μM of rFXIII-A_2_ (cell) against 25 μM rFXIII-B (syringe). We analysed the resulting isotherms using Origin 7.0 (Originlab) and fitted the data using Affinimeter and a custom model based on stepwise association of the subunits: FS ↔ MA+A1↔MA2; where FS is free species, M is FXIII-A_2_ in cell, and A is FXIII-B from syringe. To examine FXIII-A_2_B_2_ complex disassembly, we titrated 1.25 mM FXIII-A_2_B_2_ in the cell [13.8 U Thrombin (Sigma-Aldrich Chemie GmbH, Taufkirchen, Germany)] against 25 mM CaCl_2_ in the syringe. We performed blank experiments to account for the heat of dilution. We first analysed data for a single set of binding models using Origin software, to observe binding as a global fit. We then calculated heat capacity changes for each injection based on algorithms within the Origin software and stoichiometric equilibria model (described below) in Affinimeter (https://www.affinimeter.com), and iterated between these until no further significant improvement in fit was observed. Data were fit using the custom design model and hypothetical equation: M1+A1⇋M1A1+A1⇋M1A2+A1⇋M1A3; where M is FXIII-A_2_B_2_, and A is calcium ion.

## RESULTS

### AFM topographs indicates complex formation restricts the conformational flexibility of FXIII-B

AFM analysis of the FXIII-A_2_B_2_ complex revealed that each isolated surface height signal had a bi-partite appearance (*Supplemental Figure 7*) comprised of a clearly compact part (FXIII-A subunit) from which filamentous signals (FXIII-B subunit) extended in different directions (**Figure 1**). The maximum height observed in the topographic images (raw) for the FXIII-A_2_B_2_ complex was 5.9 nm for whole field, which was lower than those recorded for either FXIII-A_2_ subunit (9.5 nm) and FXIII-B_2_ subunits (19.2 nm) (**Figure 1**, *Supplemental Figure 7*). This demonstrates that the association of FXIII-B_2_ with FXIII-A_2_ restricts the conformational flexibility of free FXIII-B_2_ subunit making it more compact. In the surface topographic images only, part of the dispersed flexible region i.e. FXIII-B subunit is visible peeking out from underneath the compact part i.e. the FXIII-A subunit. This can be explained by overall negative charge carried by FXIII-A_2_ dimer surface, which relies on positive electrostatic patches on the FXIII-B subunit to adhere to the mica surface in a complexed state (**Figure 1d**). The differences in height might be attributed to adsorption effects on the structure of the protein(31, 32). Wrapping of FXIII-B subunits around FXIII-A_2_ dimer occurred from one side, giving the molecule a bi-partite appearance, suggesting partial asymmetry in the complex.

**Figure 1.**
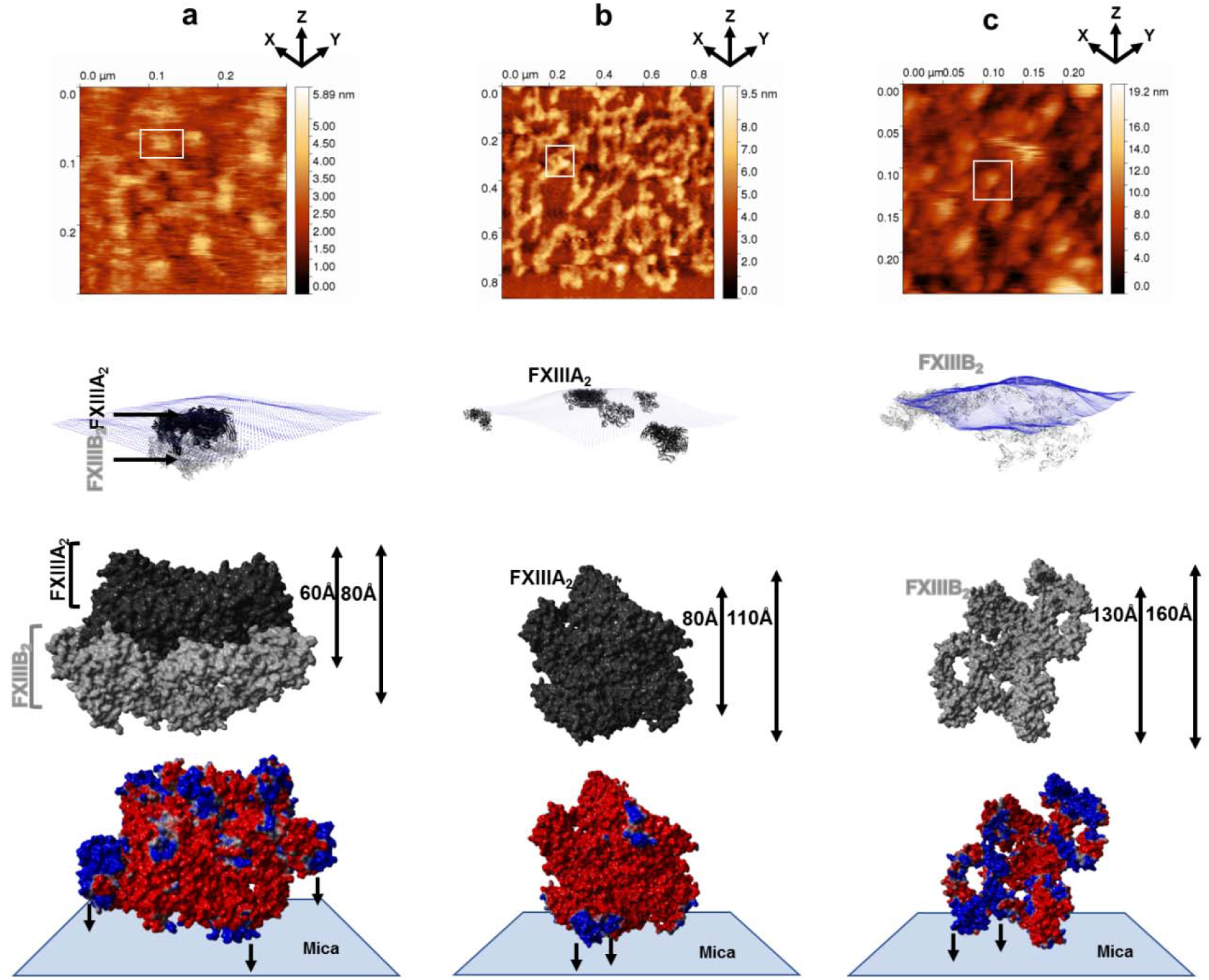
Conformational state of FXIII complex, FXIII-A, and FXIII-B subunits by AFM. This Figure is split row-wise into Panels a, b and c running top to bottom. Figure 1a goes top to bottom in the following order for the AFM and AFM based docking of the best FXIII-A_2_B_2_ complex model: the topmost image is the raw AFM image with the docking pose of one of the crops below it. In the docking pose the topography is depicted as blue dots while the different docked complexes (of model 1 only) are depicted in black (FXIII-A_2_) and gray (FXIII-B_2_) ribbon format. Below the docking pose is a molecular surface-based representation of FXIII-A_2_B_2_ complex as it would be viewed in one of the many poses it would adopt while adhering to the mica in the AFM instrument. The minimum and maximum heights that this pose is likely to have, is indicated to the right. The FXIII-A and FXIII-B subunits are depicted in black and gray color respectively. The lowermost image is PME electrostatic surface structural representation of the same pose depicted in alignment with the hypothetical mica surface to which it adheres. Figure 1b and 1c panels follow the same trend as Figure 1a, only they represent the FXIII-A dimeric crystal structure and the dimeric unbound FXIII-B model respectively.

### Crosslinks in the FXIII-A_2_B_2_ complex interface expose reverse, N-to C-terminal symmetry between FXIII-A and FXIII-B subunits

We then used XL-MS to identify inter- and intra-molecular contacts within the FXIII-A_2_B_2_ complex. This analysis generated 358 total peptides, with an overall coverage of 80% for FXIII-A and 91% for FXIII-B. The cross-linked FXIII-A_2_B_2_ heterotetramer (MW 319.950 kDa) had 34 cross-linked peptides located within the heterotetramer interface (*Supplemental Table 3*). Intersubunit crosslinks were detected between residues from the FXIII-A C-terminal barrel domains and the FXIII-B N-terminal S1, S2, and S3 sushi domains, while residues in the FXIII-A N-terminal β-sandwich domain were cross-linked to residues from the FXIII-B C-terminal S6, S7, S8 and S9 sushi domains (**Figure 2a**). The FXIII-A catalytic core region was cross-linked to FXIII-B sushi domains S3, S4, S5, S7, S8 and S9. Intra-subunit crosslinks with the FXIII-B_2_ dimer interface largely involved residues in the N-terminal sushi domains (S1-S4), but fewer cross-links within sushi domains S6, S7, and S8 (**Figure 2b**; *Supplemental Table 3*). These findings differ from those previously reported for inter- and intra-subunit interactions within the FXIII-A_2_B_2_ complex(1, 6).

**Figure 2.**
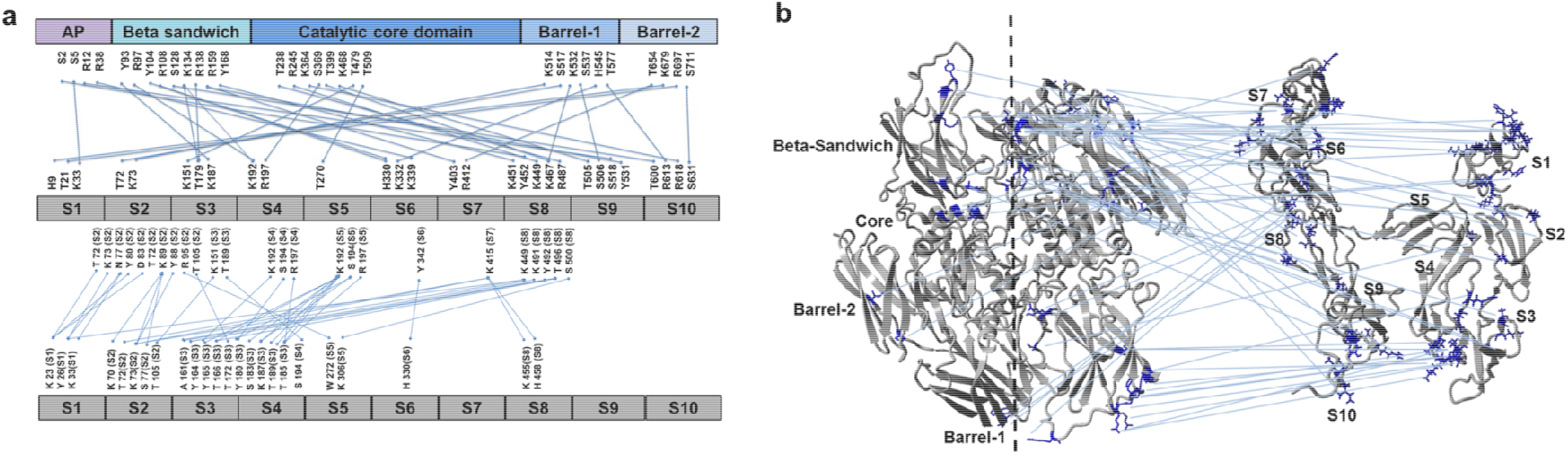
XL-MS derived crosslinking residues of FXIII complex reveals an N to C terminal symmetry. **Figure 2a** shows the domain-wise distribution of both FXIII-B monomer model docking to FXIII-A_2_ crystal structure-distance constraints (upper part of image) and monomer FXIII-B model assembly- distance constraints (lower part of image) that were generated from the XL-MS crosslinking information of the purified FXIII-A_2_B_2_ heterotetramer complex (Supplemental tables 2 and 3). **Figure 2b** a structural description of the information shown in Fig. 2a. The crystal structure of the FXIII-A subunit dimer and the monomer model of the FXIII-B subunit have been illustrated in ribbon format.

### Molecular docking reveals a stoichiometrically-symmetrical, bi-partite, FXIII-A_2_B_2_ complex

To understand the origin of the reverse symmetry of the FXIII-A_2_B_2_ complex observed in the XL-MS data, we used molecular docking to model FXIII-A_2_B_2_ assembly. Of 5 potential models of FXIII-B monomer (*Supplemental Figure 4*), only two gave successfully docked clusters with FXIII-A_2_; of these, we selected the topmost model (*Supplemental Figure 8*; chosen based on HADDOCK scores) of top-most docking cluster as our model of choice based on agreement with structural information from AFM. This model (model 1) also illustrated a symmetric bi-partite structure, in which FXIII-A subunits are compact, and FXIII-B subunits are more dispersed/flexible (**Figure 3a**). Following equilibration, backbone RMSD/total energy charts (**Figure 3b**) indicated this model of FXIII-A_2_B_2_ showed good stability and was stereochemically validated (*Supplemental Figure 9*). Discrepancies in validation were like those observed for standard complex crystal structures. The FXIII-B N-terminal sushi domains (S1, S2 and S3 domains) extended into flexible arms (**Figure 3c**), although each FXIII-B monomer showed different flexibility and secondary structure following the simulation (**Figures 3d and 3e**; *Supplemental video 1*). We observed three distinct positively-charged electrostatic patches on FXIII-B (**Figures 3f and 3g**), which may represent potential fibrinogen interaction interfaces(33) since these would create excellent complementarity with the negatively charged regions within the currently-proposed FXIII-B interaction site on fibrinogen(33–37).

**Figure 3.**
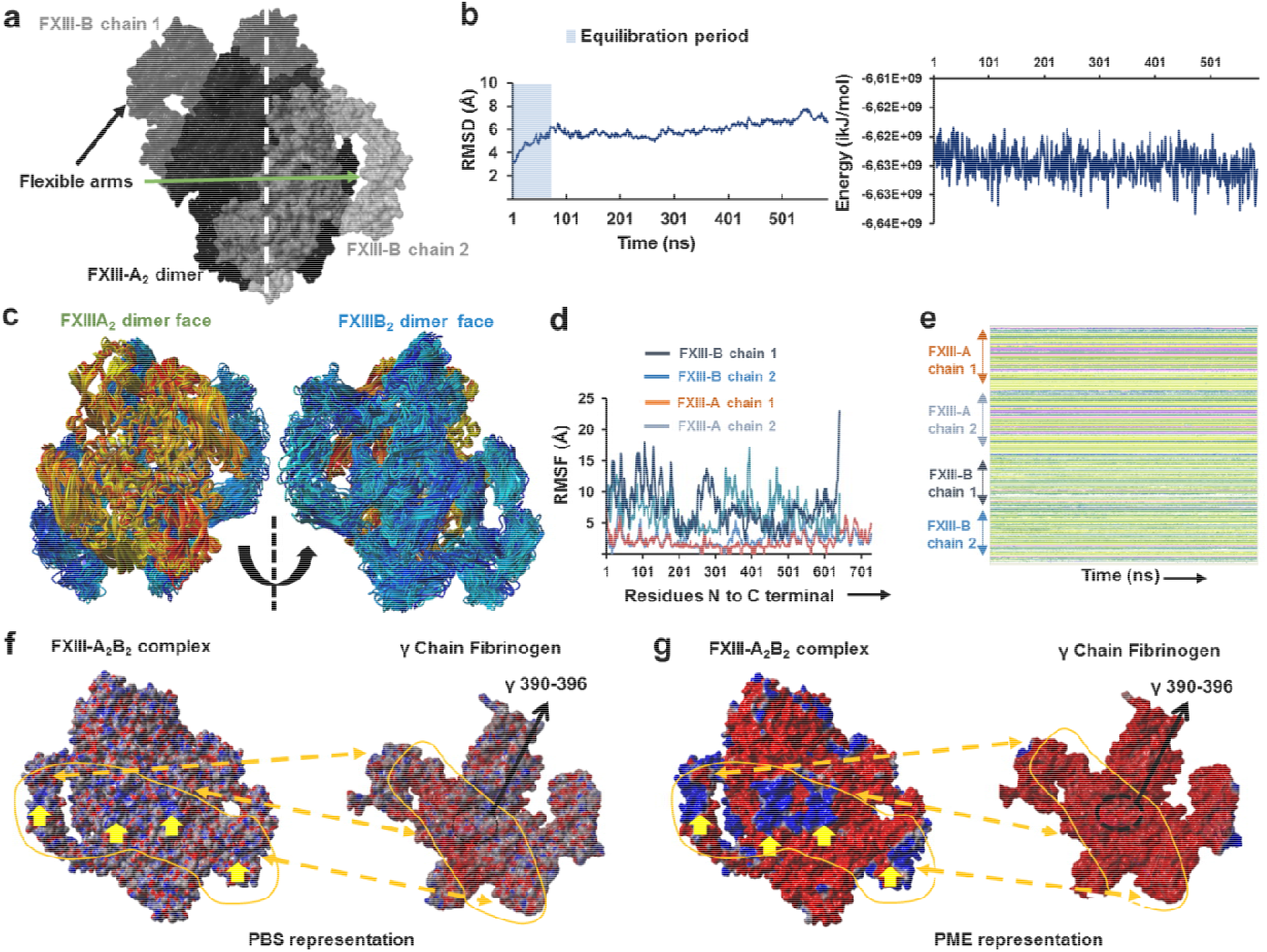
The all-atom structure of the FXIII-A_2_B_2_ complex. **Figure 3a** is the symmetrical representation of the best modeled all-atom structure of the FXIII-A_2_B_2_ complex. The structure has been depicted by its molecular surface in different shades of black and gray for the individual chains of FXIII-A/ FXIII-B subunits. **Figure 3b** shows the C-α backbone RMSD and the total energy graphs for the MD simulation conducted on the FXIII-A_2_B_2_ complex structure model. **Figure 3c** are aligned simulation snapshots from the MD simulation conducted on FXIII-A_2_B_2_ complex structure represented with FXIII-A subunit face (left) and the FXIII-B subunit face (right). The snapshots of FXIII-A/FXIII-B subunits are depicted in ribbon format with colors ranging between yellow-red and cyan-blue for either subunit respectively. **Figure 3d** shows the graph representing RMSF for the FXIII-A_2_B_2_ complex structure MD simulation; with individual chains represented by different color as mentioned in the inset; and Figure 3e represents the secondary structure profile of individual chains of FXIII-A/FXIII-B subunit for the FXIII-A_2_B_2_ complex structure MD simulation. **Figure 3f** is the PBS based electrostatic surface representation of the FXIII-A_2_B_2_ complex structure (left) and γ chain of fibrinogen (right) taken from fibrinogen crystal structure (PDB ID: 3GHG). Red color indicates negative surface electrostatic potential while blue represents positive potential. Indicated positive electrostatic patches on the FXIII-A_2_B_2_ complex structure that are likely to interact with negatively charge bearing regions in and around the FXIII interaction site of fibrinogen γ chain (the specific residues are numbered and indicated with a black arrow). **Figure 3g** is the same view as Fig. 3f but electrostatic surface representation has been done with the PME method.

### Molecular docking into the AFM topographs identifies the best model representative of the native FXIII complex

To rule out false positive conformational models of the FXIII-A_2_B_2_ complex, we docked the two FXIII-A_2_B_2_ modeled structures onto the AFM topography image. Based on the docking scores (AFM dock), FXIII-A_2_B_2_ complex model 1 had globally better scores than complex model 2 for the 10 selected docked regions of the AFM images (**Figure 4**). Each isolated surface height signal had a bi-partite appearance (*Supplemental Figure 7*) comprised of a compact part (FXIII-A_2_) from which filamentous signals (FXIII-B) extended in different directions. The AFM topographs were in line with the complex model 1 appearance. The complex model 1 therefore represents the native conformation of FXIII complex.

**Figure 4.**
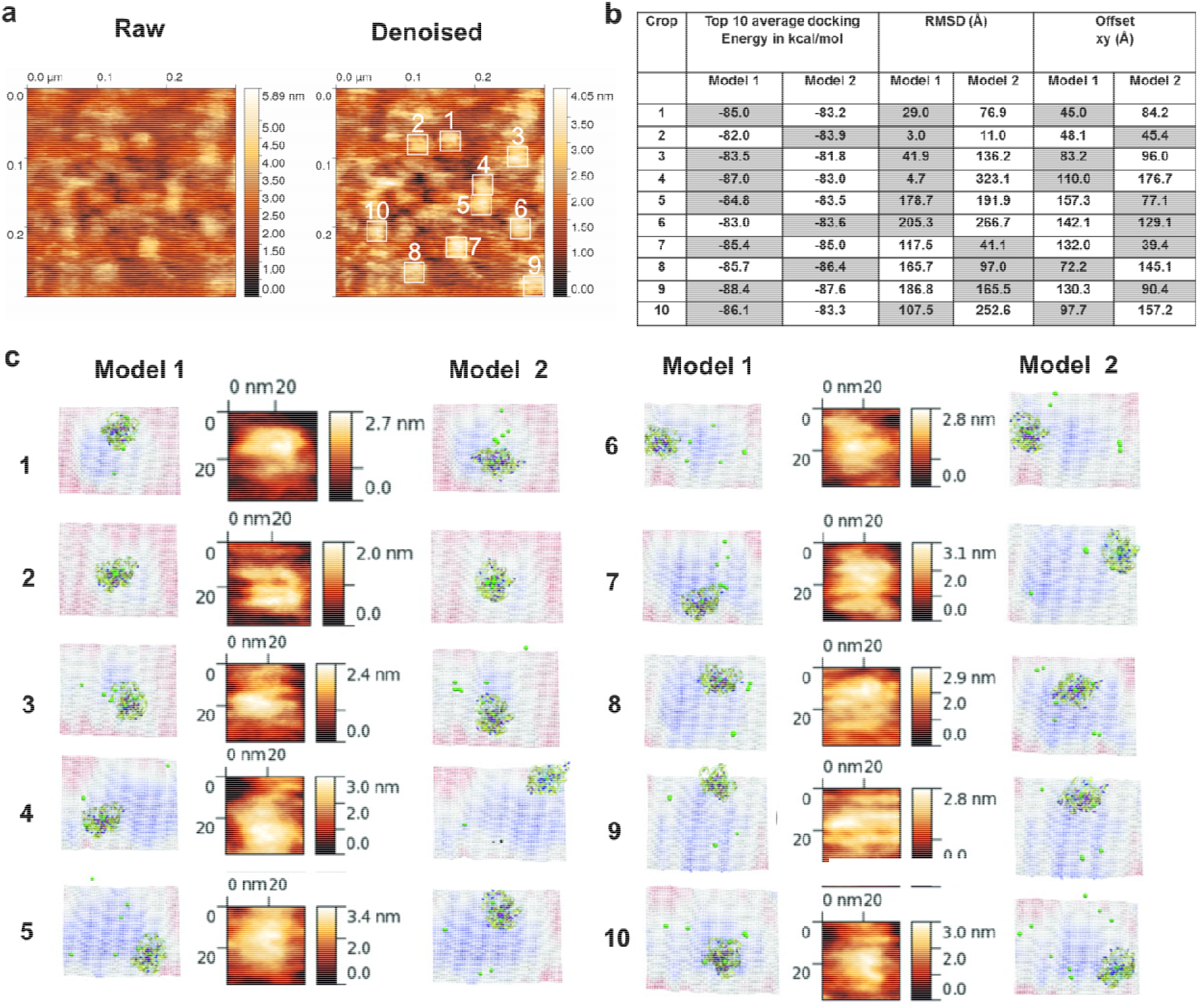
AFM based docking of FXIII complex models reveals Model 1 as best complex. **Figure 4a** shows the raw and denoised AFM topographic images for the purified FXIII-A_2_B_2_ complex. The height scales are depicted to their right. The denoised image also shows in white lined squares the crops on the topographic surface to which docking of the two best models (HADDOCK scores) of the FXIII-A_2_B_2_ complex were performed on the DockAFM pipeline(30). **Figure 4b** is a table presenting the comparative scores obtained from the docking of the two FXIII-A_2_B_2_ complex model structures on the ten AFM image crops depicted in **Figure 4a**. the xy (offset) represents the shift of the docked model structure (model 1 and model 2) from the center of the topographic surface. The most favorable structure is chosen as that having the smallest shift from the center. **Figure 4c** shows side by side the best docking pose for the two FXIII-A_2_B_2_ complex model structures on each of the ten crops side by side to a close-up topographic view of the crop itself. The topography of the docked pose is inverted i.e. looking from below the surface. The color of the topography (blue to red) is the height in Z (red is low, blue is high). The structures of the two models are depicted in ribbon format.

### Unequal pairing within the bi-partite FXIII-A_2_B_2_ complex influences dissociation of subunits during FXIII activation

Analysis of the final model 1 of the FXIII-A_2_B_2_ complex indicated inequality in binding of individual FXIII-B monomers to the FXIII-A_2_ dimer (**Figure 5a**), as well as comparative differences in pseudo-binding energies (from both simulation and PRODIGY calculations)(38) calculated for FXIII-B monomers and FXIII-A_2_ subunits (**Figures 5b and 5c**). SMD-based separation of FXIII-B_2_ from FXIII-A_2_ showed that even though steering force was only applied to FXIII-B, it displaced one of the FXIII-A monomers, resulting in a FXIII-A_2_B heterotrimer-like scenario (*Supplemental video 2*). Unequal pairing results in disproportionate displacement of FXIII-A subunits, wherein strong affinity between FXIII-A and FXIII-B subunits causes the FXIII-A subunits to be pulled apart along with the FXIII-B subunit. This observation raises the possibility that a FXIII-AB heterodimer is generated transiently during activation-induced disassembly of the complex. Thus, unequal pairing of FXIII subunits results in a previously unrecognized dissociation pattern of FXIII-B and -A subunits during FXIII activation.

**Figure 5.**
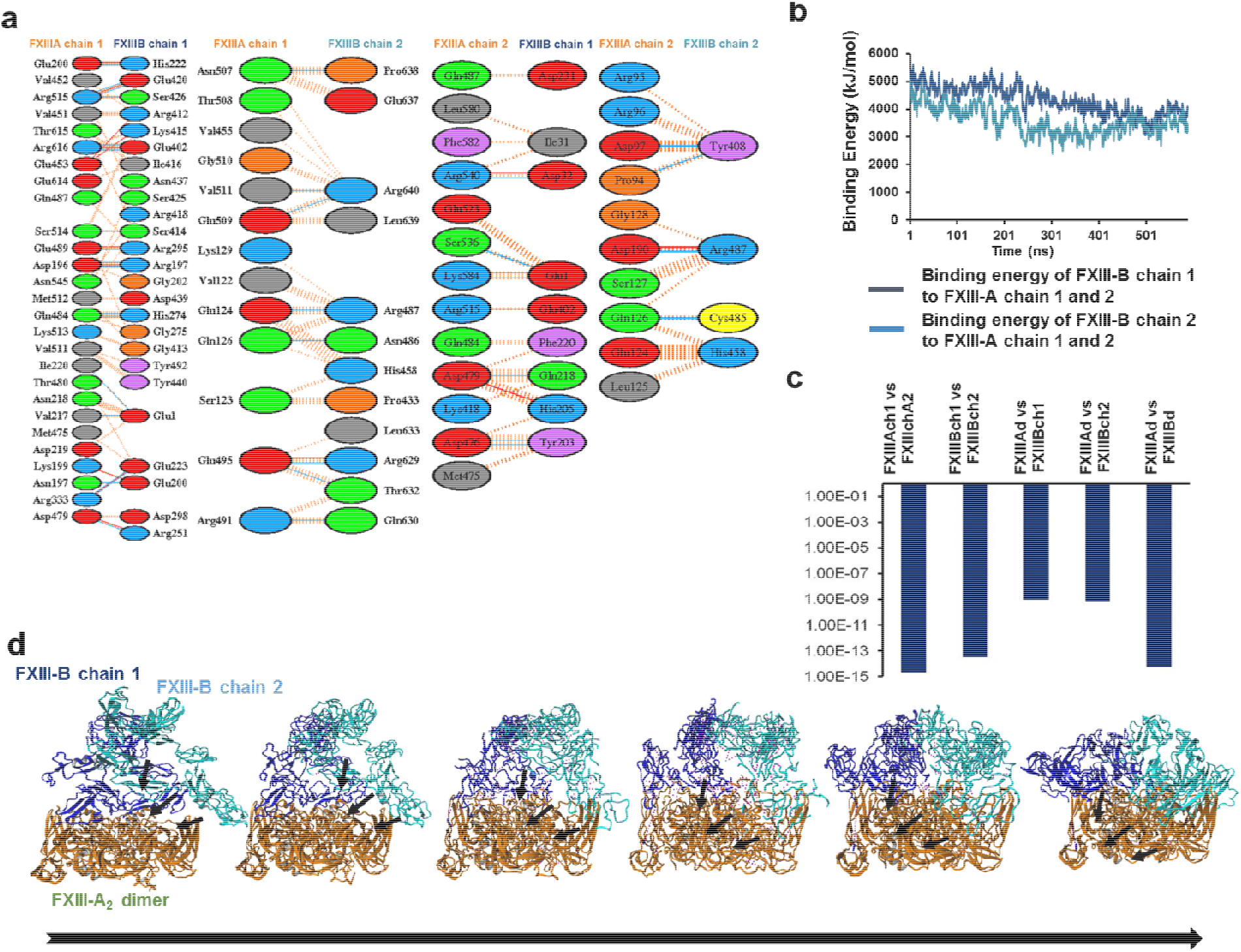
Inter-domain interactions, binding affinity and the assembly of the best FXIII-A_2_B_2_ complex model structure. **Figure 5a** shows the different type of interactions between the different chains of FXIII-A and FXIII-B subunits within the best FXIII-A_2_B_2_ complex model structure. **Figure 5b** shows a comparative binding energy graph of the two individual chains of the FXIII-B subunit to the FXIII-A_2_ dimer as calculated during the MD simulation conducted in the best FXIII-A_2_B_2_ complex model structure. **Figure 5c** show the comparative predicted binding affinities for different structural entities within the best FXIII-A_2_B_2_ complex model structure as calculated over the PRODIGY server(38). These entities are abbreviated as: FXIII-Ach1/FXIII-Ach2: FXIII-A subunit chain 1 and 2, FXIII-Bch1/FXIII-Bch2: FXIII-B subunit chain 1 and chain 2, FXIII-Ad/FXIII-Bd: FXIII-A and FXIII-B subunit dimers, respectively. **Figure 5d** shows the conformational transitions taking place in the FXIII-B dimer during its association with the FXIII-A_2_ subunit dimer. Both the subunits are depicted in ribbon format. Solid arrows represent the conformations adopted by FXIII-B during its association. The FXIII-A_2_ subunit dimer is colored orange while the individual monomers of FXIII-B subunit dimer are colored blue and cyan respectively.

### Transition state analysis suggests stepwise binding events during the association of FXIII-A: FXIII-B subunits

Since FXIII-A_2_B_2_ assembly occurs in plasma, we modeled the association events taking place during FXIII-A_2_ and FXIII-B_2_ interaction leading up to complex formation using transition state analysis. The transition state analysis of the heterotetramer assembly shows that the association of dimeric subunits to form a complex is asymmetric, two-step binding, where FXIII-B monomer first strongly associates with FXIII-A_2_ dimer, stabilizing a transient state FXIII-A_2_B’B (where B’ represents the unbound monomer). This complex then forms the FXIII-A_2_B_2_ complex where all subunits interact in totality (**Figure 5d**). These data show that the two-step asymmetrical binding most likely results in unequal pairing between the monomers within the complex.

### Thermodynamic patterns underlying FXIII-A_2_B_2_ complex assembly and dissociation suggest stepwise models for both events

Finally, having investigated the complex assembly and disassembly (during activation) events at a structural level, we performed ITC for the same events to explain the operant thermodynamic variables. ITC enabled us to a) examine the thermal changes corresponding to protein interface interactions upon binding and b) correlate the thermal motions derived from activation-induced disassembly of the complex to the structural dynamics of individual subunits obtained by the model analyses. The first set of ITC experiments were performed to illustrate the thermal mechanics underlying the binding of FXIII subunits. When fitted with one set of binding sites, data measuring association of FXIII-A_2_ and FXIII-B_2_ yielded a K_d_ of 66.7 nM. A sequential 2-step binding model based on hints from the unequally paired FXIII-A_2_B_2_ complex model 1 suggested that the first binding event of FXIII-B_2_ to FXIII-A_2_ is an enthalpically-favorable exothermic reaction (ΔH= −226.25 kJ/mol), yet yields a conformationally-restricted state with positive, unfavorable entropy (-TΔS>0) and K_d_ of 1.5 nM. The second binding event (MA↔MA_2_) is also enthalpically favorable (ΔH= −360 kJ/mol) with unfavorable entropic changes (-TΔS>0), but a comparably weaker K_d_ of 4.3 μM, due to the spatial restriction faced by the second monomer upon interaction (**Figure 6**). These thermodynamic patterns agree with our transition state analysis suggesting two-step binding assembly, the latter being of low affinity, leading to heterotetramer assembly.

**Figure 6.**
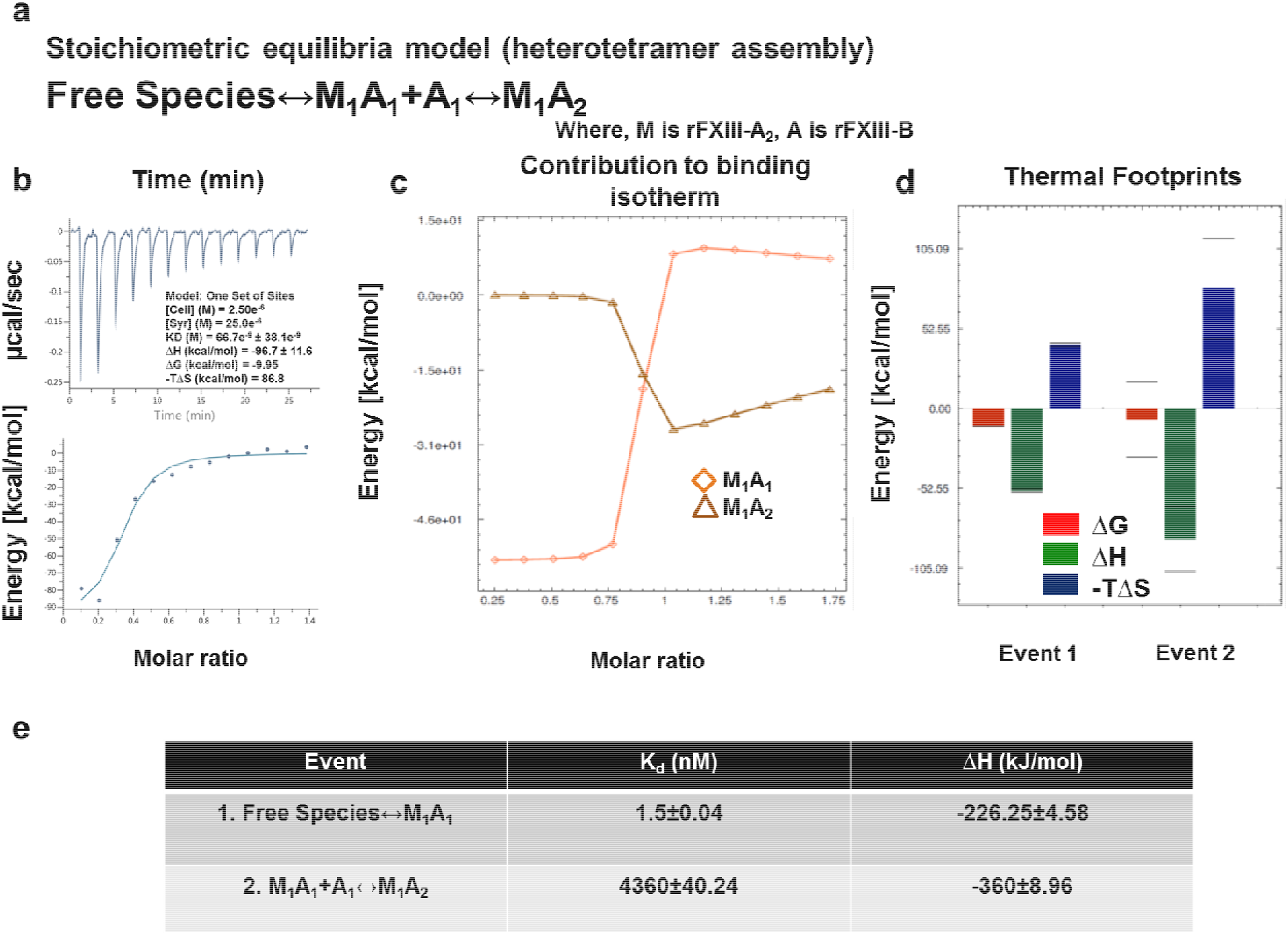
In-solution associations of FXIII subunits studied by ITC. **Figure 6a** is the equation depicting the stoichiometric binding equilibrium model; followed for the analysis of data derived from ITC (Model was generated in Affinimeter using model builder approach). **Figure 6b** represents the titration of 2.5μM rFXIII-A_2_ (in cell), with 25μM FXIII-B subunits (in syringe). Upper image of this panel is the raw data depicting the heat change upon each injection; lower image in this panel is the normalized data, with integrated heat change plotted against the concentration ratio of rFXIII-B vs rFXIII-A_2_. (Blank controls not shown) Solid black line represents the corresponding fit obtained in Origin software using one-set of binding mode. **Figures 6c** and **6d** are based on Affinimeter analyses depicting the contribution of individual reactants of equation (Fig. 6a) towards the isotherm; the heat signatures depicting the free energy changes, changes in enthalpy and entropy in the two events explained in Fig. 6a respectively. **Figure 6e** is table explaining the two thermodynamic events, and their corresponding Dissociation constants (K_d_) and changes in Enthalpy (ΔH).

To analyze the dissociation of the complex upon activation, we also interrogated calcium binding to the thrombin-cleaved FXIII-A_2_B_2_ complex. Based on the stoichiometric equilibrium model (**Figure 7a**) post-fitting, the first calcium binding event (set at K_d_ 100 μM(39–41)) showed an entropically-driven, negative -TΔS, and unfavorable, endothermic ΔH (4.78 kJ/mol) pattern. The second event, corresponding to K_d_ of 1 mM, had highly negative -TΔS, and endothermic ΔH (150.70 kJ/mol) behavior. In contrast, the third event, corresponding to K_d_ of 1.94 μM, had a highly positive -TΔS and exothermic ΔH (−154.42 kJ/mol) (**Figure 7**) heat change. Experiments assessing thrombin- and calcium-mediated dissociation of FXIII-A_2_B_2_ suggested the events proceeded step-wise. (i) Calcium binds to FXIII-A_2_ in the heterotetramer complex. (ii) The calcium-bound heterodimer separates (i.e. FXIII-AB). Given heat signatures (**Figure 7d**) obtained for event 2 (ΔH>0, -TΔS>0), we propose that the system maintains thermal equilibrium by first dissociating into a transient FXIII-AB heterodimer. This step combats the unfavorable enthalpy of A/A or B/B subunits suggested by the *in silico* pseudo-binding energy calculated for our complex model 1 (**Figure 5c**). (iii) The FXIII-AB heterodimer separates into individual, free subunits. During this event, unfavorable conformational entropies are counteracted by favorable enthalpic changes, which explain the final disruption of FXIII-AB heterodimer into calcium saturated, activated and open FXIII-A* monomer(31)). At the conditions used (T=30°C), all three events were spontaneous (ΔG<0). The flipping patterns of enthalpy and entropy in the subevents that occur during dissociation of the complex suggests the role of bulk solvent coming into play, along with calcium saturation of FXIII-A, that is responsible for stepwise disassembly of complex(42, 43)’(44). Collectively, our thermodynamic data support the premise that FXIII undergoes two unique stepwise modes of complex assembly and disassembly in plasma (*Supplemental Figure 10*).

**Figure 7.**
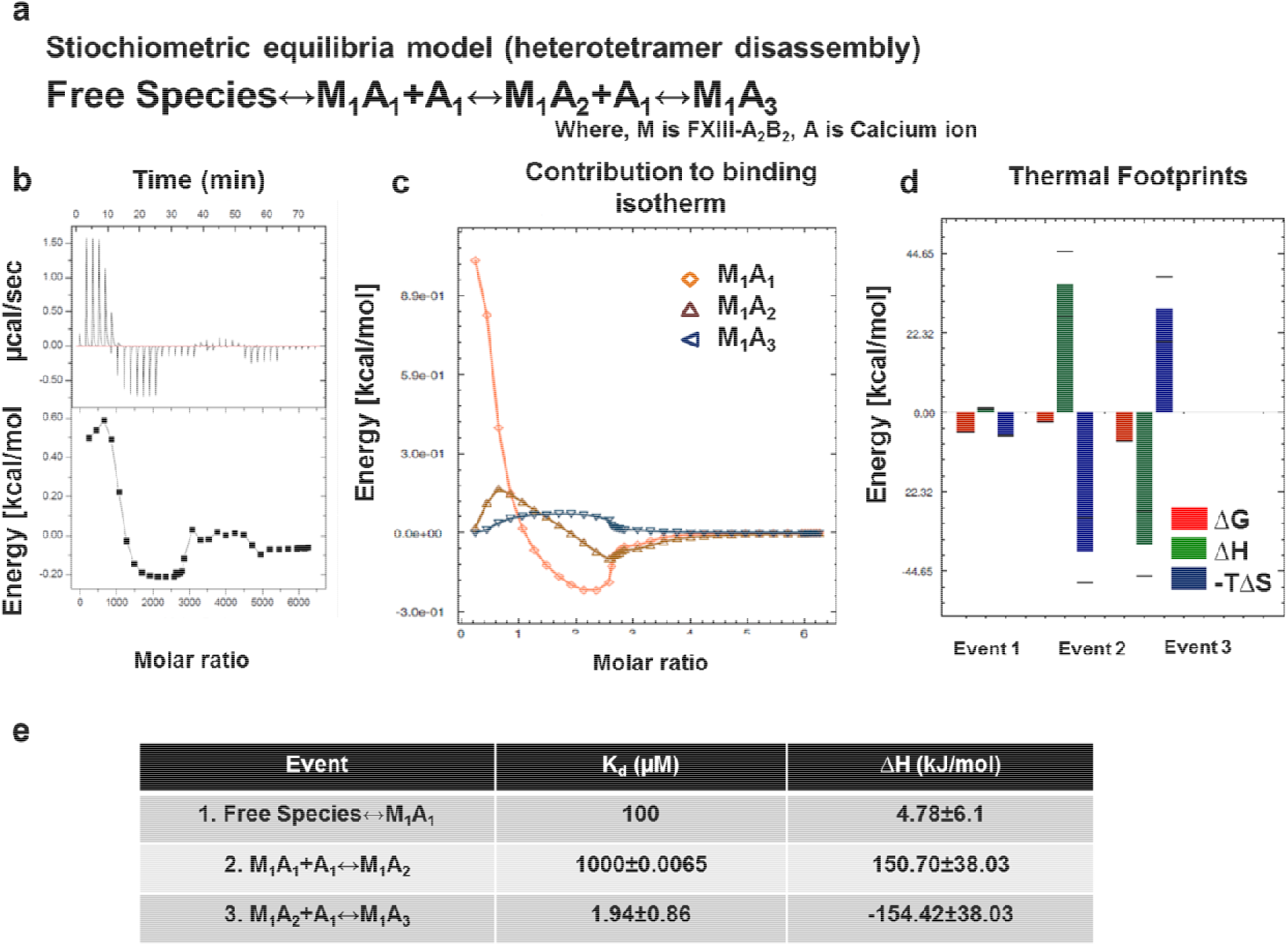
In-solution dissociation of FXIII complex in the presence of thrombin and calcium studied by ITC. **Figure 7a** is the equation depicting the stoichiometric binding equilibrium model; followed for the analysis of data derived from ITC (Model was generated in Affinimeter using model builder approach). **Figure 7b** represents the titration of 1.25mM FXIII-A_2_B_2_, with 25mM CaCl_2_. Upper image of this panel is the ORIGIN raw data depicting the heat change upon each injection; lower image in this panel is the normalized data, with integrated heat change plotted against the concentration ratio of CaCl_2_ vs FXIII. Solid black line represents the corresponding fit obtained in Origin software using one-set of binding mode. (Blank controls not shown) **Figures 7c and 7d** are based on Affinimeter analyses depicting the contribution of individual reactants of equation (Fig. 7a) towards the isotherm; the heat signatures depicting the free energy changes, changes in enthalpy and entropy in the two events explained in Fig. 6a respectively. **Figure 7e** is table explaining the three thermodynamic events, and their corresponding Dissociation constants (K_d_) and changes in Enthalpy (ΔH).

## DISCUSSION

### IH approaches reveal a unique FXIII complex structure

The FXIII complex has always presented a structural functional challenge to researchers owing to its dynamic nature and association with various other proteins like fibrinogen in its physiological and biochemical life-cycle. The absence of an all atom basis for this complex further limits interpretation of pathomolecular mechanisms, as well as the development of drugs/inhibitors to bind both complexed and isolated FXIII-A(45–48). Indirect evidence in the context of inter-domain interactions exists, but does not have a visual/structural basis(1). The final structure of the FXIII-A_2_B_2_ complex model derived in this study fills this gap. Notably, our model presents a new picture of the complex that differs from the current paradigm in several important ways^1,6^. First, our model suggests FXIII-B subunit N-terminal sushi domains are relatively free, with positive surface electrostatic patches, indicating a potential role of this flexible region in interactions with other proteins like fibrinogen^30,31,33^. These interactions would be especially relevant when evaluating the effect of FXIII or fibrinogen surface mutations that disrupt or disorder their mutual complex. In addition, reports have suggested the N-terminal region of FXIII-B interacts with the FXIII-A subunit(1). This observation could also be a secondary allosteric effect observed during the competitive binding studies performed; since both our all atom structure as well as the XL-MS analysis suggest increased density of interdomain interactions at the C-terminal end of FXIII-B. The recent report by Proptopopova et al(49), based primarily on AFM studies in air suggests partial wrapping of B subunits around central core of FXIII-A_2_. Our study registers similar observations, but we propose a different orientation possibility for individual subunits within the complex. Differences between our interpretations might be due to the surface properties of the a) complex, b) the HOPG/mica surface, c) sample preparation, or d) imaging in air or buffer, which affects the overall behavior of the molecule under the microscope(32, 50). Association analyses by ITC reveals that the binding of individual subunits is strong (~10^−9^ M; single set of binding mode) but differs by a factor of 10 from the latest report (~10^−10^ M) in which one of the subunits was immobilized, unlike our solution based label free evaluation(1).

### Assembly of FXIII-A_2_B_2_ is a two-step process aided by the conformational flexibility of FXIII-B subunit

Assembly of the FXIII-A_2_B_2_ complex in plasma and its subsequent activation and disassembly are well known phenomenon. However, the thermodynamic variables underlying these events have not been investigated in detail, especially in the context of the conformational changes that occur during these processes. Published data address only the thermolability of individual domains of FXIII-A subunit(51). In contrast to transglutaminase-2 (TG2), a near homologue of FXIII-A which has been thoroughly investigated in the thermodynamic aspect(52), detailed data for the FXIII complex is not available. Our ITC experiments performed in-solution balance our structural investigation into FXIII oligomeric association and disassembly upon activation by identifying subtle changes in thermodynamic parameters that fit into our structural models. The thermodynamic study of FXIII complex assembly was performed by titrating both subunits against each other. Step-wise association of the two FXIII-B subunits onto FXIII-A dimer, also depicted by the *in-silico* transition state models, suggests the conformation adopted by the transient transition state FXIII-A_2_BB’ supports final complex association (**Figure 5d**). The heat signatures observed in our study show that desolvation of the inter-subunit interface leading to the stabilization of non-covalent contacts across the interface is a gradual two-step event aided by the conformational flexibility of FXIII-B subunits. While the globular FXIII-A structure remains relatively static, the flexible FXIII-B monomers twist and bend to individually accommodate the FXIII-A monomers. This two-step, unequal, asymmetrical assembly, further supported by the significantly different binding energies for the two events observed in our ITC profile (**Figure 6**), is the underlying cause for an unequally paired complex. This analysis is additionally supported by our AFM-based observations, in which the free form of FXIII-B is flexible and long but once associated with the FXIII-A, becomes more compact. Regardless of this association, even complex-bound FXIII-B retains some flexibility, especially at its N-terminal regions as observed from the thermal motions of the complex model simulation (*Supplemental video 1*). These observations also suggest the FXIII-B subunit undergoes a significant conformational change during its transition from a free molecule to a bound one.

### Unequal pairing within the FXIII complex may generate a transient FXIII-AB species during the activation induced complex disassembly

We studied FXIII complex disassembly by saturating thrombin-cleaved FXIII-A_2_B_2_ with increasing concentrations of calcium in an ITC platform. The thermodynamic driving forces responsible for complex activation indicate the role of solvation energies(43) that release the subunits in the presence of calcium ions (**Figure 7**). Here the transition of the zymogenic heterotetramer to active, open, monomeric FXIII-A* involves formation of a transient FXIII-AB heterodimer in which FXIII-A subunits are incompletely saturated with calcium and are still loosely bound to one of the FXIII-B monomers. Our thermodynamic analyses indicate how low-entropy interfacial water molecules(42, 53, 54) assist in disrupting the tetrameric interface in FXIII-A_2_B_2_, aiding the entropic compensations that act against unfavorable enthalpies. Relevance of this model of dissociation becomes clearer when examining the nature of unequal pairing within the complex. Unequal pairing of individual FXIII-B monomers, especially at the C-terminal hinge region of FXIII-A (residues 500-520) (*Supplemental Figure 11a*; *Supplemental videos 3 and 4*), is favorable since it enables bulk solvent to sneak past the interface between loosely-bound B monomer and A subunits (*Supplemental Figure 11b*), aiding dissociation to perform timed activation. This flexible hinge region is critical for movement of FXIII-A barrel-domains, enabling its activation and giving rise to its open extended conformation(8). Equally strong binding of both FXIII-B subunit monomers to FXIII-A_2_ would be energetically expensive, also yielding possibly slower activation/subunit dissociation. However, this observation does not suggest the complex breaks down into a trimer (FXIII-A_2_B) and a monomer (FXIII-B), as is also observed in our SMD for separation of FXIII-B_2_ from FXIII-A_2_. Notably, bulk solvent/ water does not play a role during the separation in the SMD (*Supplemental video 2*), unlike a physiological scenario in which it is an active participant. Bulk solvent/water permeates interfaces between all subunits, bringing a semblance of symmetry to the disruption/disassembly process in the physiological environment. We can, however, conclude that the process of dissociation is strong enough to separate FXIII-A monomers from each other (*Supplemental video 2*; *Supplemental Figure 12*), providing further support that the activated FXIII-A molecule is a monomer(31). Given the binding affinity of the FXIII subunits and their conformational motions, FXIII mutations that affect interface residues or conformational flexibility are likely to undermine complex assembly, resulting in either loosely or too tightly bound complex. A tightly-bound complex will trap the oligomer in less flexible states and alter rates of activation, whereas a loosely-held complex may be susceptible to spontaneous disassembly. The models of assembly and disassembly during activation of FXIII complex implicated by our study are particularly relevant in the context of the structural data and thermodynamics for research pharmacologists interested in generating inhibitors/drugs directed against the complex. The steps detailed in these events especially in the thermodynamic context can be objectively addressed to virtually and actually screen for inhibitors as has been done for TG2, FXIII’s homologue(55).

### Is complex interface a potential underlying driver of unexplained heterozygous FXIII mutations observed in mild FXIII deficiency?

The catalytic FXIII-A subunit bears a special place in the transglutaminase (TGase) family as it is the only member that exists in a complexed form (FXIII-A_2_B_2_). Consequently, interfacial residues within the complex are under selective pressure, wherein mutations at these residues might be associated with broad range of factor deficit (mild to severe FXIII deficiency). Inspection of recently reported missense mutations of *F13A1* genes (p.His342Tyr, p.Asp405His, p.Gly411Cys, p.Gln416Arg, p.Leu539Pro, p.Arg540Gln, p.Gln601Lys, p.Arg611His)(56–59) suggests these mutations lie on the interface rim(60) where they may be involved in FXIII-A and FXIII-B subunit interactions. Similarly, FXIII-B missense mutations (p.Cys336Phe, p.Val401Glu, p.Pro428Ser and p.Cys430Phe)(61, 62), also map to interfacial patches. Notably, a majority of these mutations were reported in the heterozygous state from mild FXIII deficiency patients. Since inherited FXIII deficiency is an autosomal recessive disorder, the dominant negative effect of heterozygous FXIII gene mutations might be explained by their pathomolecular influence on the complex interface.

### Implications for coagulation biochemistry and therapeutic potential

To summarize, our study presents not only a new view of the FXIII complex, but also proposes new mechanisms to explain FXIII complex association and disassembly. The structure provides a basis on which FXIII mutations (particularly those thought to affect the FXIII molecular interface) can be probed to define their pathomolecular mechanisms. Furthermore, our model provides the first atomic basis on which putative inhibitors can be designed and tested. Our models present interesting starting points for research into conformational changes occurring during FXIII complex assembly and disassembly. Further biochemical validation of hypotheses stemming from these models is warranted.

## Authorship Contributions

A.B. conceived and designed the project. S.S. performed the protein expression, purification, and ITC. A.N. performed and analyzed XL-MS. S.S and S.K. performed AFM. J-L.P. performed AFM dock analyses. A.B. performed all the in-silico analyses. S.S. and A.B., analyzed the data and co-wrote the manuscript. A.S.W, D.I. and J.O. read and edited the manuscript. All authors critically reviewed the manuscript.

