## Supplementary information for "The plasma Factor XIII heterotetrameric complex structure: unexpected unequal pairing within a symmetric complex"

***^b^*** *CovalX, Schützengasse 2, CH-8001 Zürich, Switzerland*

***^c^*** *German Center for Neurodegenerative Diseases (DZNE), Sigmund-Freud-Str. 27, 53127, Bonn, Germany*

***^d^*** *University of Grenoble Alpes, CEA, CNRS, IBS, F-38000, Grenoble, France*

***^e^*** *Department of Pathology and Laboratory Medicine, University of North Carolina at Chapel Hill, 8018A Mary Ellen Jones Building, Chapel Hill, North Carolina 27599-7035, United States*

***^f^*** *Pharmaceutical Biochemistry and Bioanalytics, Pharmaceutical Institute, University of Bonn, An der Immenburg 4, 53121 Bonn, Germany*

**Supplementary materials and methods**

***FXIII-A subunit, cloning expression and purification in Yeast based Pichia Pink expression system***

Cloning expression and purification of rFXIII-A (recombinant FXIII-A) subunit was performed as described by Gupta et al, 2016^1^. Briefly, the gene of interest was flanked with a combination of Tags (10X His, Strep, and FLAG) separated by TEV protease cleavage site. Aforementioned construct was expressed in *Pichia pastoris* host strains (*PichiaPink*™ strain 3 ade2-/-, prb1-/-; Invitrogen, Germany) in BMGY media, induced by methanol in BMMY media for 48 hours. After cell harvesting, a two-step purification approach was employed to purify the full-length rFXIII-A protein based on His-tag, using His-TRAP column (GE Healthcare, Sweden), followed by a second purification based on immuno-affinity for streptactin tag (Iba Lifesciences, Germany). Elute containing rFXIII-A was further purified by gel filtration using a Superdex 200 Increase 3.2/300 (GE Healthcare, Germany) size exclusion chromatography (SEC) column until a single, homogenous, mono-dispersed peak was obtained. The protein was resuspended in 20mM Tris-Cl, 120mM NaCl, 1mM EDTA (pH 7.4) buffer.

***FXIII-B subunit, expression and purification in HEK293T mammalian expression cells***

The human *HEK293t* cell line purchased from DMSZ (German Collection of Microorganisms and Cell Cultures) were cultured in high glucose DMEM (Invitrogen, Germany) supplemented with 10% v/v FBS (Invitrogen, Germany), 1% v/v Penicillin-Streptomycin antibiotics (Invitrogen, Germany) and 0.1% v/v Fungisone (Invitrogen, Germany), at 37°C at 5% CO_2_. All experiments were done on sub-cultured cells in logarithmic phase (below passage 20). Human FXIII-B cDNA, inserted into the cloning site of pReciever-M01 mammalian expression vector was transfected into *HEK293t* cells as per previously reported protocol^2^. Briefly, 2.7 X 10^5^ cells were transfected with 3 µg of plasmid DNA along with 6 µl of transfection reagent lipofectamine 2000 (Invitrogen, Germany). Cultures were harvested 48 hours post-transfection, by collecting extracellular fractions containing the secreted rFXIII-B (recombinant FXIII-B). The extracellular fraction was centrifuged for 5 minutes, at 14000 rpm at 4°C. A negative control of non-transfected cells was used, whereas a plasmid construct with eGFP cloned in pcDNA mammalian expression vector, was the positive control for transfection. Secreted protein harvested post transfection of *HEK293t* cells was concentrated and subjected to immuno-affinity based purification using the Thermo Scientific Pierce Co-IP kit (Pierce Biotechnology, USA) following manufacturer’s protocol. Briefly, monoclonal antibodies against FXIII-B, raised in mice were immobilized to Amino Link plus coupling resin (Pierce Biotechnology, USA) for 2 hours. The resin was then washed with PBS and incubated with extracellular concentrate overnight in cold-room. Next day, resin was again washed with PBS and protein bound to anti-FXIII-B antibodies was eluted with acidic elution buffer provided with the kit. Eluted protein was verified on coomassie stained gel. Eluted protein was further subjected to gel filtration chromatography, to ensure purity and dimeric state of the recombinant protein.

**Purification of FXIII-A_2_B_2_ from plasma concentrate FibrogamminP**

FibrogamminP (CSL Behring, Germany), was reconstituted in water (as per manufacturer’s instructions). The reconstituted FibrogamminP was purified by SEC using Superdex 200 increase 10/300 column (GE healthcare, Germany) on Äkta Pure system equilibrated with 20mM Tris, 100mM NaCl at pH 7.4. Peak corresponding to FXIII-A_2_B_2_ (320Kda), was re-purified thrice until a highly pure, single, homogenous, monodispersed peak was obtained with no excipients. The eluted peak was concentrated and quantified for further downstream applications.

**XL-MS based characterization of FXIII-A_2_B_2_ heterotetramer complex**

The purified complex FXIII-A_2_B_2_ was first analyzed for sample quality, and stoichiometry by high mass MALDI-MS analyses. Ten microliters of 400µg/ml purified FXIII-A_2_B_2_ was serially diluted to 3.12 µg/ml concentration. One microliter of each dilution was mixed with 1 μl of a matrix composed of a re-crystallized sinapinic acid matrix (10 mg/ml) in acetonitrile/water (1:1, v/v), TFA 0.1% (K200 MALDI Kit; CovalX, Switzerland). After mixing, 1μl of each sample was spotted on the MALDI plate. After crystallization at room temperature, the plate was introduced in the MALDI mass spectrometer (Ultraflex III MALDI ToF , Bruker, Germany) equipped with HM2 High-Mass detection (CovalX, Switzerland) and analyzed immediately in High-mass MALDI mode. The sample mixtures were also used for cross-linking experiments. Nine microliters of mixture was mixed with 1 µL of DSS stabilizer reagent (2mg/mL) and incubated at room temperature for 3 hours, using an earlier reported protocol^3^. The instrument was operated in linear and positive mode by applying an accelerating voltage of 20 kV and using a delay extraction of 400 ns. The HM2 system was operated in the high mass mode with acceleration voltage set to 20 kV and gain voltage set to 3.14 kV. For each sample, three spots were analyzed. The MS data were analyzed using Complex Tracker analysis software (CovalX, Switzerland). For characterization and peptide mass fingerprinting, the purified FXIII-A_2_B_2_ complex was submitted to ASP-N, trypsin, chymotrypsin, elastase and thermolysin proteolysis, followed by nLC-LTQ Orbitrap MS/MS analysis (ASP-N: 1:200, overnight at 37°C, trypsin and elastase: 1:100, overnight at 37°C; chymotrypsin: 1:200, overnight at 25°C, thermolysin: 1:50, overnight at 70°C. After digestion, formic acid 1% final was added to the solution). The purified FXIII-A_2_B_2_ (1.25µM) was crosslinked with 2 µL of DSS reagent (d0d12 reagent (Creative Molecules Inc, Canada) at room temperature for 3 hours, prior to digestion. A nano-LC chromatography was performed using an Ultimate 3000 (Dionex, USA) system in line with an LTQ Orbitrap XL mass spectrometer (ThermoFischer Scientific, USA). After proteolysis 10µL of digested sample was loaded onto a nano-liquid chromatography system (Ultimate 3000, Dionex, USA). Briefly, peptides were cleaned and concentrated in a C4 PepMap LC column (ThermoFischer Scientific, USA). After drying, the peptides were resuspended in 5% ACN and 0.1% formic acid solution. 10µl of peptide mixture were loaded onto C4PepMap column (ThermoFischer Scientific, USA) (pre-column 300-μm ID x 5-mm C4 PepMap, pre-column flow rate 30 μl/min, column 75-μm ID x 5-cm C4 PepMapTM, column flow rate 200 nl/min). Peptides were eluted during a 5-40% gradient (buffer A: 5% ACN/0.1% formic acid, buffer B: 20% ACN/0.1% formic acid), over 35 minutes and detected in LTQ-Orbitrap (ThermoFischer Scientific, USA). The general MS conditions were: needle voltage 1.8 V; capillary voltage 5 V. The Orbitrap was operated with full-scan spectra over the m/z range 300-1700, acquired in a positive ion mode. The MS method used was double play with CID activation; the normalized collision energy was 35; the activation time was 30 ms; and the activation Q value was 0.25. Acquired data were analyzed by XQuest version 2.0 and Stavrox version 2.1. The FXIII-B intra-subunit and FXIII-A-FXIII-B inter-subunit cross-linked peptides and residues are presented in Supplementary Tables 2 and 3.

**Generation of FXIII-B subunit monomer model**

The XL-MS crosslinks were categorized into three groups: I) cross-links within the same subunit monomer II) within the same subunit but different monomers (both intra-subunit) and III) between two different subunits (inter-subunit). We first used the information contained within the FXIII-B subunit intra-subunit XL-MS cross-links to assemble a constrained monomeric model of FXIII-B subunit. We used 5 different inter-residues contact data prediction tools and servers based on different mathematical algorithms^4–8^. For each of the predictions made on individual servers we compared them to the XL-MS crosslink data and chose those that matched, to further use as constraints for assembling the monomer model of FXIII-B subunit (Supplementary Tables 4). The chosen constraints were not necessarily the exact cross-linked residues rather these were residues that were close to both the predicted residue as well as the XL-MS cross-linked residue. The monomer FXIII-B subunit was assembled from high quality threaded models of sushi domains generated on ITASSER on the AIDA domain joining and minimizing server (<http://aida.godziklab.org/>)^9^ using the XL-MS crosslinks matched to the inter-residue contact prediction data as a constraint. As a result of different prediction tools, we had 5 monomeric model conformations, all of which were candidates for the fold the FXIII-B subunit assumes in its bound form (as FXIII-A_2_B_2_). These 5 models were used to dock onto the FXIII-A_2_ crystal structure (PDB ID:1f13) to generate the heterotetramer complex models and further downstream processing. In addition, we also assembled a FXIII-B subunit monomer model on the AIDA server in default mode i.e. without any constraint (Supplementary Figure 5). This model which is expected to represent the unbound FXIII-B (because it has not been generated with XL-MS cross-linking constraints from the heterotetramer) was symmetrically docked on the MZ-docking server^10^ and the highest scoring dimer with interactions between the sushi domains S4 and S9 (based on earlier reported evidence^11^) was chosen as the final model of the unbound FXIII-B_2_ dimer.

**Constrained docking of the FXIII-B subunit model on the FXIII-A_2_ crystal structure to generate the FXIII-A_2_B_2_ all atom complex model/ structure**

Docking was performed based on the assumptions; i) the heterotetramer is a symmetrical 2:2 structure, ii) all the subunits have contact/interaction with each other within the complex. A constrained docking on the HADDOCK expert interface webserver for flexible docking was performed based on the inter-subunit crosslinks between the FXIII-A and FXIII-B subunits generated by XL-MS analyses. Since this webserver allows for only bi-molecular docking and the proteins involved comprised of three molecules, i.e. one subunit of the FXIII-B subunit monomer model and two subunits of the crystal structure of FXIII-A_2_ (PDB ID: 1f13), we treated the dimeric crystal structure as a single subunit i.e. we renumbered the residues of one of the monomer to appear as a continuation of the other monomer and also matched the XL-MS cross-linked residue data on the FXIII-B subunit similarly. This strategy was based on an initial set of calculations that we performed for the spatial positioning of FXIII-B subunit model on a monomer of FXIII-A molecule (also taken from PBD ID: 1f13) on the DisVis webserver (<http://milou.science.uu.nl/cgi/services/DISVIS/disvis/>)^12^. The constraints chosen from the XL-MS inter-subunit crosslinked data, were not necessarily the exact cross-links rather set of residues close to the actual cross-links and belonging to the same cross-linked peptide but capable of forming side chain contacts (Supplementary Table 4). We observed that utilizing even ≥12 XL-MS crosslinking directed constraints yielded no complex structure that could satisfy all the constraints spatially. Therefore, in order to cover the entire surface of the FXIII-A_2_ subunit dimer we used 64 XL-MS cross-linking directed constraints for the flexible docking of the FXIII-B subunit monomer model on the crystal structure of the FXIII-A_2_ dimer instead of on a FXIII-A monomer. It should be kept in mind that the PDB file of FXIII-A_2_ dimer i.e. 1f13 has missing atoms and loops. These were modeled on the FREAD loop modeling server (<http://opig.stats.ox.ac.uk/webapps/fread/>)^13^ and the final gap resolved structure was used for docking purposes. The distances for the constrained docking assigned between the FXIII-A and FXIII-B residues were kept at a constant 24 Å keeping in mind the length of the cross-linker (DSS) used as well the flexibility of side chains^14^.

**Simulation of the heterotetramer FXIII-A_2_B_2_ models and other *in silico* assessments**

The stability of the top-scoring FXIII-A_2_B_2_ complexes (Supplementary Figure 1) generated from the HADDOCK server was analyzed by running all atom molecular dynamics (MD) simulation on the YASARA Structure package version 13.11.1 platform^15^. The PDB files were initially subjected to a 500 ps refinement MD simulation run that imposes the YAMBER3 force field parameters in YASARA in order to remove steric clashes and improve rotamer geometry^16^. The file with the lowest energy in the simulation trajectory was chosen for conducting further simulations. Simulations were performed with the md_sim macro embedded in YASARA. Briefly, a simulation cell with periodic boundaries and 20 Å minimum distances to protein atoms was employed with explicit solvent. The AMBER03 force field, NPT ensemble was used with long-range PME potential and a cut-off of 7.86Å^17^. The simulation cell was filled with water at a density of 0.997 g/mL and a maximum sum of all bumps per water of 1.0 Å. The entire system was energy minimized by the steepest descent to remove conformation stress within the structure, followed by simulated annealing minimization until convergence was achieved. The MD simulation was performed at 298 K. Simulations for all models were run for at least >100 ns post equilibration. Secondary structure content during the MD simulations was visualized using the md_analyzesecstr macro output embedded in YASARA on R. The pseudo-binding energies during the simulation between the individual monomeric subunits were calculated using the md_analyzebindingenergy macro embedded in YASARA. Electrostatic surface potential was calculated and graphically depicted using the Adaptive Poisson-Boltzmann Solver (PBS) integrated within YASARA^18^. The trajectory was first loaded into VMD (version 1.9.3) in the Gromacs .xtc format to obtain secondary structure predictions. Secondary structure prediction was made using the STRIDE algorithm available in VMD^19^. The 2D heat map representation of the secondary structure prediction was produced via the *Timeline* plugin (version 2.3) in VMD^19,20^. Structural image visualization, analysis and rendering were done with YASARA 13.11.1 and Chimera version 1.10.2. The final equilibrated heterotetrameric complex structure(s) were validated stereo-chemically on several model quality detection servers (5 different servers were tested by submitting the structures on a single platform of SAVES version 5.0 http://servicesn.mbi.ucla.edu/SAVES/). These structures were analyzed for inter-subunit contacts on the Profunc webserver (<http://www.ebi.ac.uk/thornton-srv/databases/ProFunc/>)^21^ and also the binding affinities between the FXIII-A and FXIII-B subunits in these structures were predicted on the Prodigy webserver (<https://nestor.science.uu.nl/prodigy/>)^22^. Later a steered molecular dynamics (SMD) simulation was performed on the MD equilibrated heterotetramer complex model 1 for dissociating the FXIII-B subunit dimer component from the FXIII-A subunit dimer. The SMD was performed with md_runsteeredmd macro embedded in YASARA with minor modifications in the steering force. Briefly, a steering force with an acceleration of 100 pm/ps^2^ was applied to displace FXIII-B subunit dimer only. The direction of the force applied was from the geometric center of the receptor to the geometric center of the ligand. No part of the whole system was restrained during the SMD to allow freedom of movement for residues being influenced by the separation of the monomers. The SMD was analyzed to the point at which no intermolecular contacts were observed between the FXIII-A monomers and/or the atoms started crossing the periodic boundaries.

**Generation of structural transition path between the first contacts between FXIII-A: FXIII-B subunits to the final consolidation of the heterotetrameric complex**

This part of the analysis was purely hypothetical and was performed primarily out of curiosity to observe how the overall complex might assemble in a step-wise manner. To do this we had to first generate a possible model of the initial contact between FXIII-A_2_ and FXIII-B_2_ dimers. We used the zymogenic crystal structure of FXIII-A_2_ dimer as used before and a dimeric model of unbound FXIII-B_2_ (generated as described earlier in methods) for docking on the Z-dock rigid docking server^23^ under default conditions. The highest scoring complex resulting from this docking attempt was considered as the first or initial contact structural state. This complex structure and the best model of the full heterotetrameric complex generated on HADDOCK^15^ were submitted to the MINACTION path server (<http://lorentz.dynstr.pasteur.fr/suny/submit.php?id0=minactionpath#submit>) as initial and final states in order to generate the putative C_α_-backbone models of the structural intermediates between these two states^24^. Once a large number of coarse-grained intermediates were generated, eight were finally selected which differed from each other by at least >2 Å RMSD (of the alpha carbons of the backbone trace). The side chains of these intermediates were modeled using the PD2 ca2main server (<http://www.sbg.bio.ic.ac.uk/~phyre2/PD2_ca2main>) to generate the corresponding full atom models^25^. Full atom models were then subjected to a structure refinement protocol as described before. Lowest energy models were selected from the simulation trajectory and analyzed.

**Atomic force microscopy of FXIII-A_2_B_2_, FXIII-A_2_ and FXIII-B_2_**

The purified FXIII-A_2_B_2_, rFXIII-A_2_ and rFXIII-B_2_ were analyzed for surface topology individually by using high-resolution AFM. Samples suspended in PBS pH 7.4 (1–2 mM) were placed on a freshly cleaved mica sheet and allowed to adsorb for 15 minutes. The non-adherent proteins were removed by washing twice with imaging buffer (10 mM Tris-HCl; 50 mM KCl; adjusted to pH 7.4). Imaging was done in oscillation mode in liquid (imaging buffer) as described earlier^26^. The images were acquired using the Nanoscope III microscope with the resolution of 512 by 512 pixels. The acquired images were processed by the built-in Nanoscope III software.

**Atomic fitting/docking protein structures within AFM surface topographs**

Docking of a three-dimensional protein coordinates below an AFM topographic surface (called envelope) has been performed with the AFM-Assembly protocol^27,28^. The original AFM height image (300 nm² and 512 px²) was flattened using Gwyddion^29^ and sent to the destripe server for the removal of vertical stripe noise^30^. Crops of the de-noised image were adjusted using Gwyddion, leading to 10 sub-images of 64 px² and 37.5 nm² (5.9 Å/px). These sub-images were selected around the maximum height of pixels from the original image and only pixels above a threshold of 8 Å were kept as the docking envelope. The docking search engine is the program DOT 2.0 embedded within the DockAFM process^28^. The docking space was made as a cubic grid of 128 x 128 x 128 grid points whose respective spacing was 3 x 3 x 2 (x, y, z, in Å). The docking envelop was located at the grid center with its base plane parallel to the X-Y plane of the grid. The favorable and forbidden layers were set to 12 Å and 20 Å, respectively. The favorable layer corresponds to the region beneath the topographic envelope where the protein structure will be docked whereas the forbidden layer corresponds to the area above that envelope. Rotations of the protein structure were set with an angular step of 6° (54,000 different rotations per grid point). A docking score was defined as the number of atoms from the protein structure located within the favorable layer and translated into pseudo-energy values where the best score corresponds to the lowest energy. The docking protocol was run on the two complex models as well as on the crystal structure of FXIII-A and the dimer models of unbound FXIII-B. For each model, only the top 10^5^ of the potential solutions were retained and only the top 10 were further analyzed. Result analyses include the minimum docking energy, the average energy of the top 10 docking solutions, the root-mean-square deviation (RMSD) of the top 3 docking solutions, and the shift of the best docking solution from the center of the docking grid.

**ITC based thermodynamic profile of the assembly and dissociation of FXIII heterotetramer complex**

Isothermal titration calorimetry experiments were carried out on a MicroCal200 microcalorimeter (Malvern Panalytics, UK). The reference cell was filled with Autoclaved MiliQ water (Millipore, USA). For examining the binding of FXIII subunits, 2.5µM rFXIII-A_2_ (purified in-house), was titrated against 25µM rFXIII-B subunit (purified in-house). The titration involved 13 injections of 3µl each. Resulting isotherm was first analyzed on Origin Software provided with the MicroCal. For custom based analyses, keeping in view the stepwise association of subunits (FS ↔ MA+A1↔MA2; where FS is free species, M is FXIII-A_2_ in cell, and A is FXIII-B from syringe), Affinimeter was used for model fitting. All the experiments were performed in the same buffer (20mM Tris, pH 8.2), at 30 degrees with the stirring speed set to 750rpm, low feedback. The final data was interpret based on fitting exercises performed using Affinimeter, using K_d_1 as the K_d_ obtained from origin analyses using one set of binding sites as a non-varying parameter.

For studying the complex disassembly, FXIII-A_2_B_2_ (Purified from FibrogamminP (CSL Behring)) in the sample cell, was titrated against calcium concentrations (in the syringe). The sample was re-suspended in 20mM Tris pH 8.2 before titration. For analyses of the thermodynamics of calcium binding, 1.25mM of FXIII-A_2_B_2_ (with 13.8U Thrombin (Sigma) in the sample cell); were titrated against 25mM CaCl_2_. The titration involved 19 injections (12×2, 0.4×6, each injection spaced by 120s). To ensure complete saturation, the reaction was continued with further 19 injections (12×2, 0.4×6, each injection spaced by 120s). Both the resulting isotherms were concatenated with CONCAT tool provided with the Origin software (version 7.0) (OriginLab). For the FXIII-A_2_B_2_ titration, thrombin was added just prior to the start of reaction, hence FXIII-A_2_B_2_ was incubated with Thrombin, in the absence of calcium for the period of pre-titration delay. To account for the heat of dilution, we performed blank experiments under the same conditions without FXIII-A_2_B_2_ in the sample cell. Peak integration was done in the software Origin 7.0. For thermodynamic analysis, initially we used Origin software, for single set of binding model, in order to observe the binding as a global fit. The parameter K_d_ , for binding of calcium to FXIII-A was set to fit around 100 µM based on earlier reports^31,32^, was set as non-varying parameter for sequential binding site model with 3 set of binding sites. (The K_1_ was used as a standalone non-varying parameter for further fitting of data in all fitting exercises). Subsequently, heat capacity changes for each injection were calculated based on the algorithms followed by Origin software, as well as stoichiometric equilibria model in Affinimeter (<https://www.affinimeter.com>) and the process was iterated until no further significant improvement in fit were observed. Individual heat-signatures were derived from the energy definitions upon each binding after model fitting also by using Affinimeter. The corresponding thermodynamic parameters were calculated according to the equation ln (1/K_d_) = (∆H-T∆S)/RT. The titrations were first simulated in Affinimeter and fitting was simulated to the following custom design model, using the model-builder approach on Affinimeter app in order to dissect the contribution of individual species generated during the course of FXIII-A activation, following the hypothetical equation: M1+A1⇋M1A1+A1⇋M1A2+A1⇋M1A3. Where, M is FXIII-A_2_B_2_, A is calcium ion.

**Supplementary Table 1** An accumulated list of software and databases used in our IH approach to characterize the FXIII complex

| **Purpose** | **Software/Webserver/Database** | **Reference** |
| --- | --- | --- |
| FXIII-B sushi domain definitions  Threaded modeling sushi domains  Submission of sushi domain models  Assembling of sushi domains  Model stereo chemical quality check  Modeling missing residues/loops  Interaction map | Uniprot (<https://www.uniprot.org/>)  ITASSER (<https://zhanglab.ccmb.med.umich.edu/I-TASSER/>)  Model Archive (https://www.modelarchKlicken Sie hier, um Text einzugeben.ive.org/)  AIDA (<http://aida.godziklab.org/>)  SAVES (<http://servicesn.mbi.ucla.edu/SAVES/>)  FREAD loop modelling server (<http://opig.stats.ox.ac.uk/webapps/fread/>)  ProFunc ( <https://www.ebi.ac.uk/thornton-srv/databases/profunc/>) | ^33^  ^34,35^  ^36,37^  ^9^  ^6,38–41^  ^13^  ^21,42^ |
| Contact residue prediction servers | Coinfold/RaptorX-Contact (<http://raptorx2.uchicago.edu/ContactMap/>)  PconsC3 (<http://pconsc3.bioinfo.se/pred/>)  ICOS (<http://cruncher.ncl.ac.uk/psp/prediction/action/home>)  SVMSEQ (<https://zhanglab.ccmb.med.umich.edu/SVMSEQ/>)  CORNET (<http://gpcr.biocomp.unibo.it/cgi/predictors/cornet/pred_cmapcgi.cgi>) | ^43^  ^5^  ^44^  ^45^  ^46^ |
| Flexible docking  Binding affinity prediction  Accessible interaction space evaluation  Multimeric rigid fast docking  Rigid fast docking | HADDOCK expert interface (<http://milou.science.uu.nl/services/HADDOCK2.2/haddockserver-expert.html>)  Prodigy (<https://nestor.science.uu.nl/prodigy/>)  DISVIS (<http://milou.science.uu.nl/cgi/services/DISVIS/disvis/>)  MZ-dock (<http://zdock.umassmed.edu/m-zdock/>)  Z-dock (<http://zdock.umassmed.edu/>) | ^15,47,48^  ^49^  ^12,50^  ^51^  ^23^ |
| Molecular dynamic simulation  Transition state models | YASARA macros  MINACTION path server (http://lorentz.dynstr.pKlicken Sie hier, um Text einzugeben.asteur.fr/suny/submit.php?id0=minactionpath#submit | ^17,18^  ^24^ |
| Mass spectrometry analysis | XQuest version 2.0  Stavrox 2.1  Complex Tracker | ^52^  ^53^ |
| Structural image/ Simulation variable visualization and depiction | YASARA, VMD, Chimera | ^17,18,54,55^ |
| AFM  AFM docking | Gwyddion  DeStripe (<http://biodev.cea.fr/destripe>)  DockAFM pipeline (<http://biodev.cea.fr/dockafm/>) | ^29^ ^30^ ^28^ |
| ITC data analysis | Origin  AFFINImeter (<https://www.affinimeter.com/site/>) | ^56,^  ^57,58^ |

**Supplementary Table 2** The FXIII-B subunit intra-subunit XL-MS cross-linked peptides and the specific residue cross-linked within the peptide.

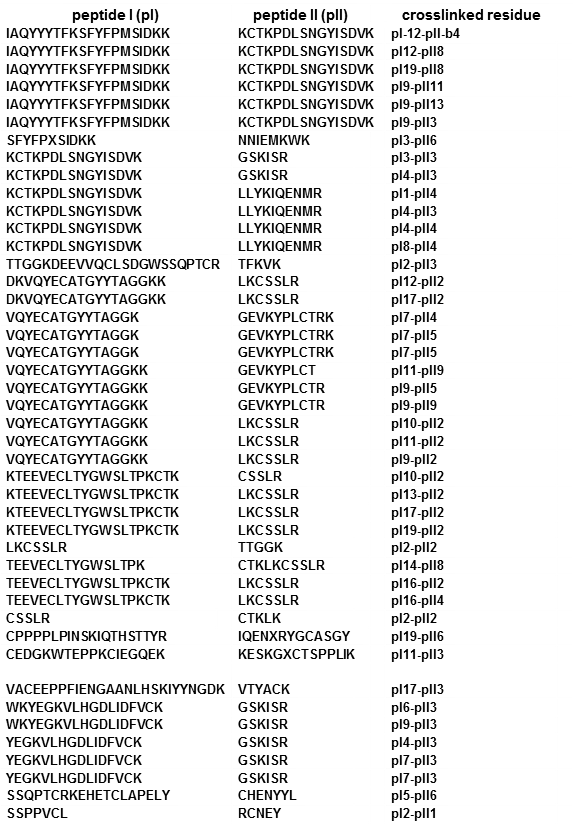

**Supplementary Table 3** The FXIII-A-FXIII-B subunit inter-subunit XL-MS cross-linked peptides and the residues cross-linked within the peptide.

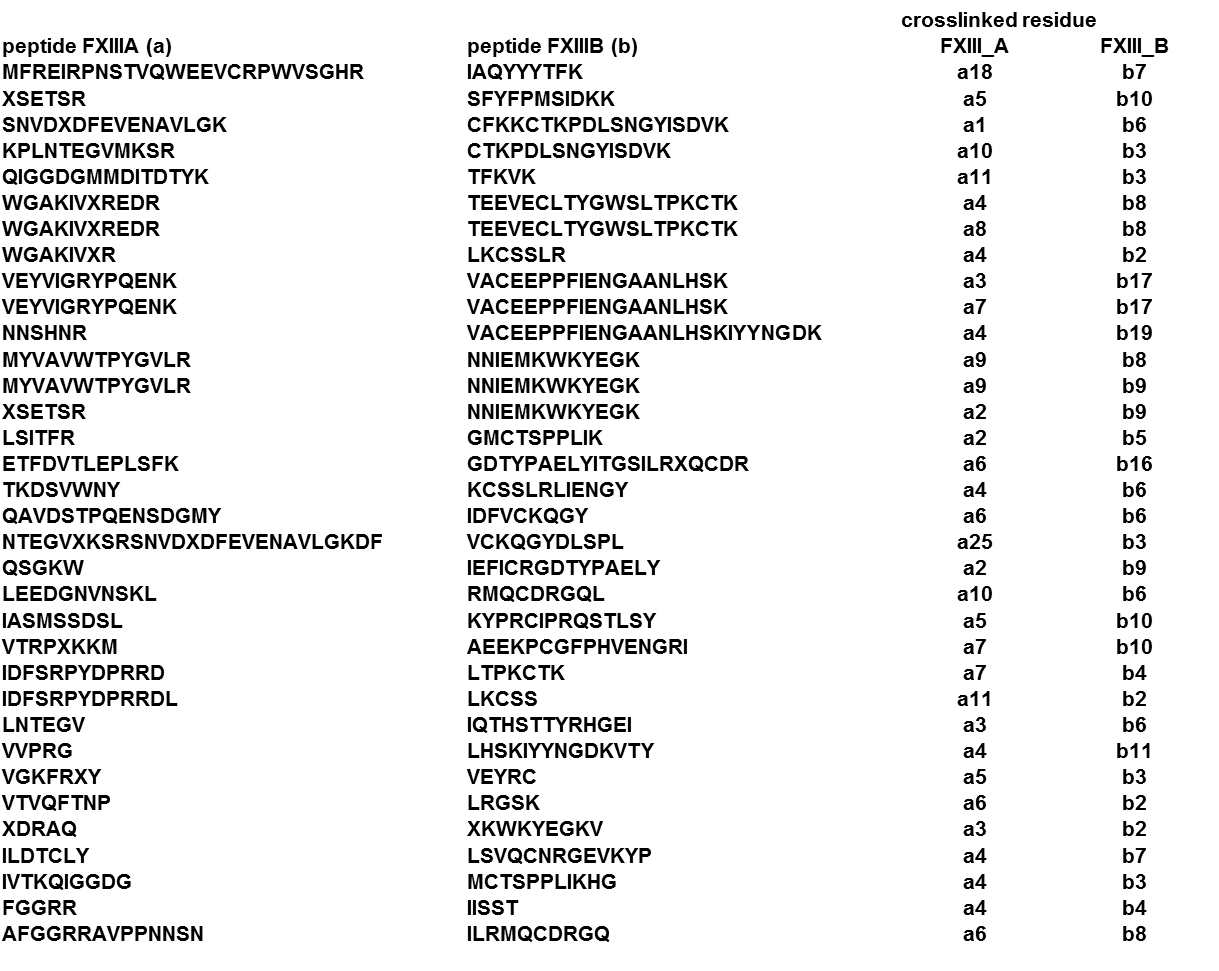

**Supplementary Table 4** The constraints built around on XL-MS data that were applied to the flexible docking of FXIII-B monomeric model on the dimeric crystal structure of FXIII-A subunit. The constraints are presented in typical HADDOCK 2.2 webserver format.

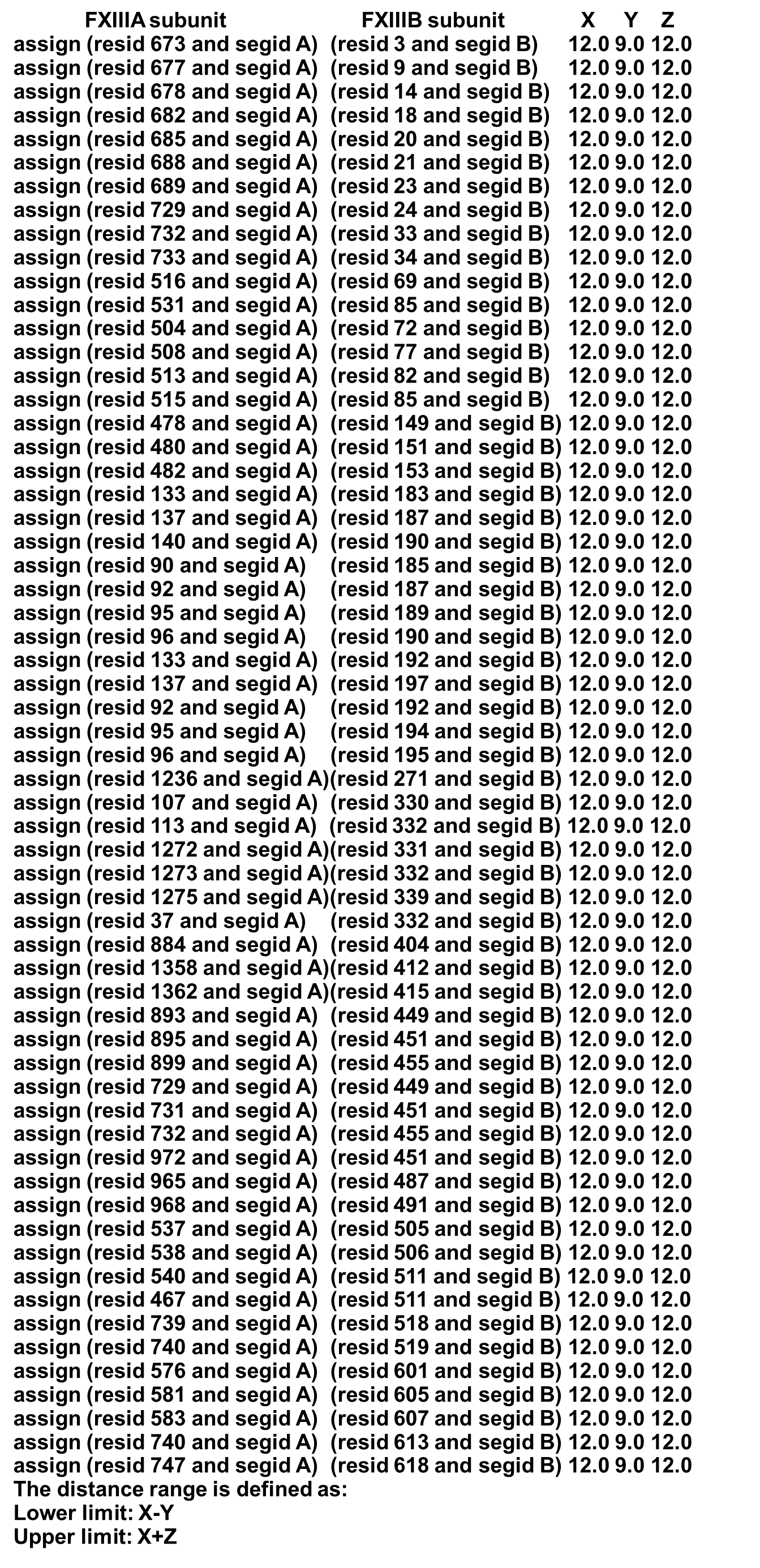

**Supplementary Figure 1** A flowchart for IH approach employed in this study to characterize the FXIII complex

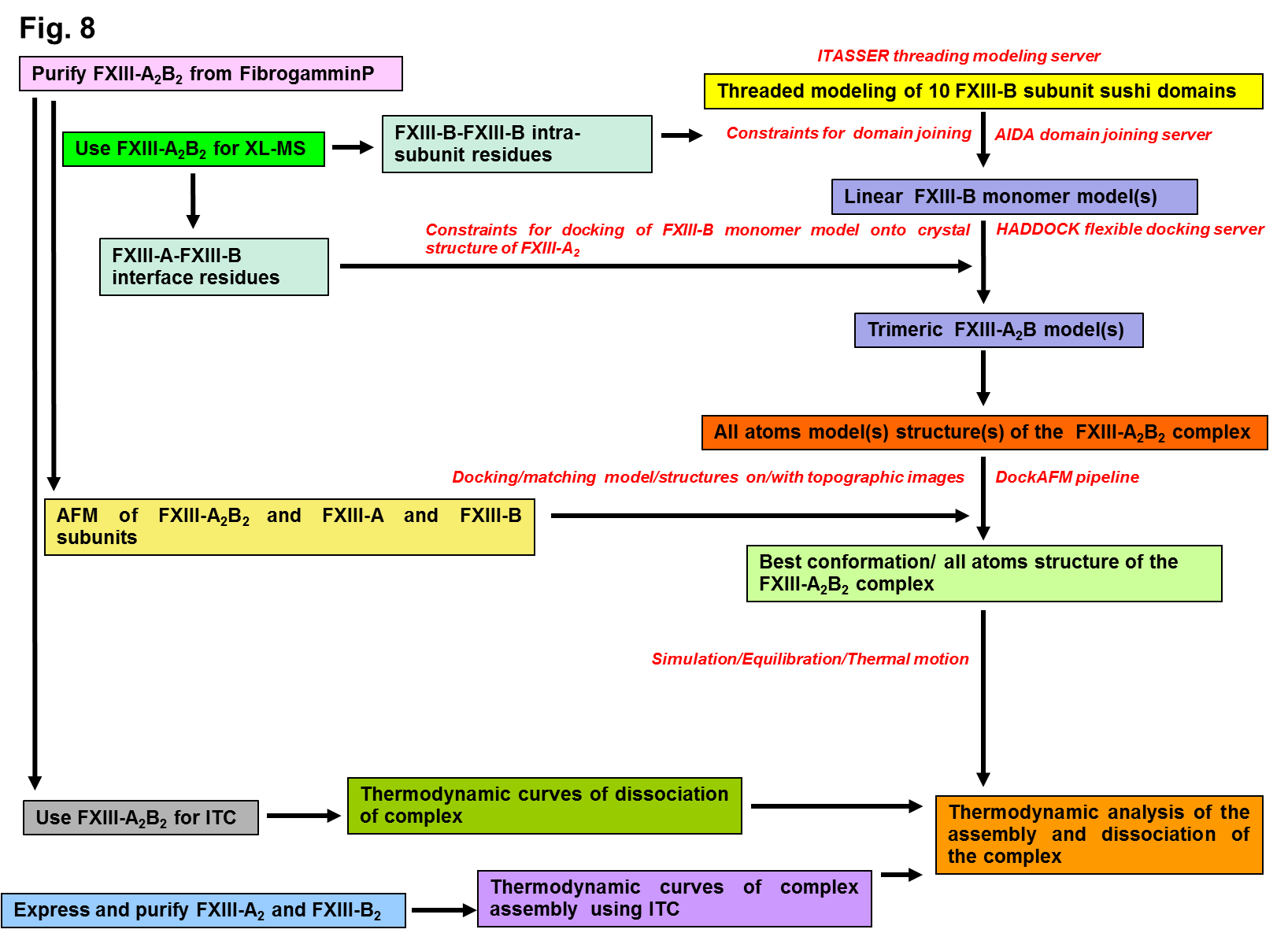

**Supplementary Figure 2a** The expression and purification of rFXIII-A subunit from *Pichia pastoris* (yeast) cells. **Supplementary Figure 2b** The expressed and purified rFXIII-B subunit from mammalian *HEK293t* cells. **Supplementary Figure 2c** The FXIII complex separated and purified from the plasma FibrogamminP concentrate.

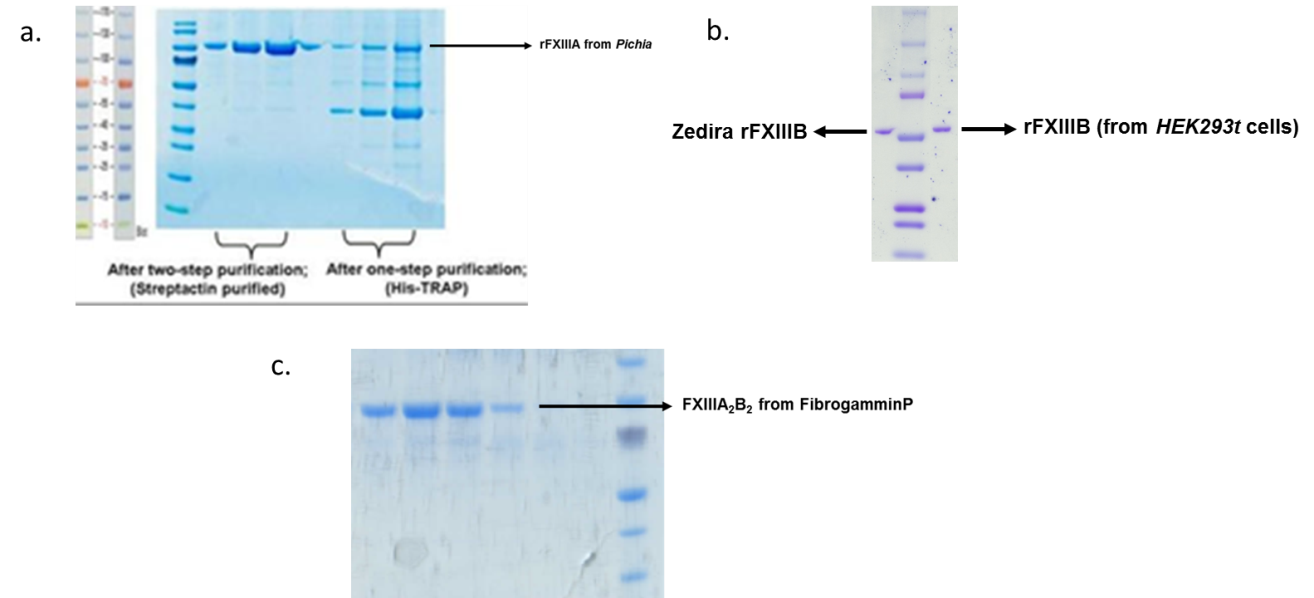

**Supplementary Figure 3** **Panel a** shows the contact residue prediction map generated from the Raptor X-contact webserver that was matched with the intra-subunit crosslinking data in Supplementary Table 2 to generate the FXIII-B subunit model domain assembly constraints shown in **Panel b**. This particular combination of constraints was applied on the AIDA server to generate the Model 1 of FXIII-B monomer which eventually resulted in the complex model 1 of the heterotetramer complex post docking.

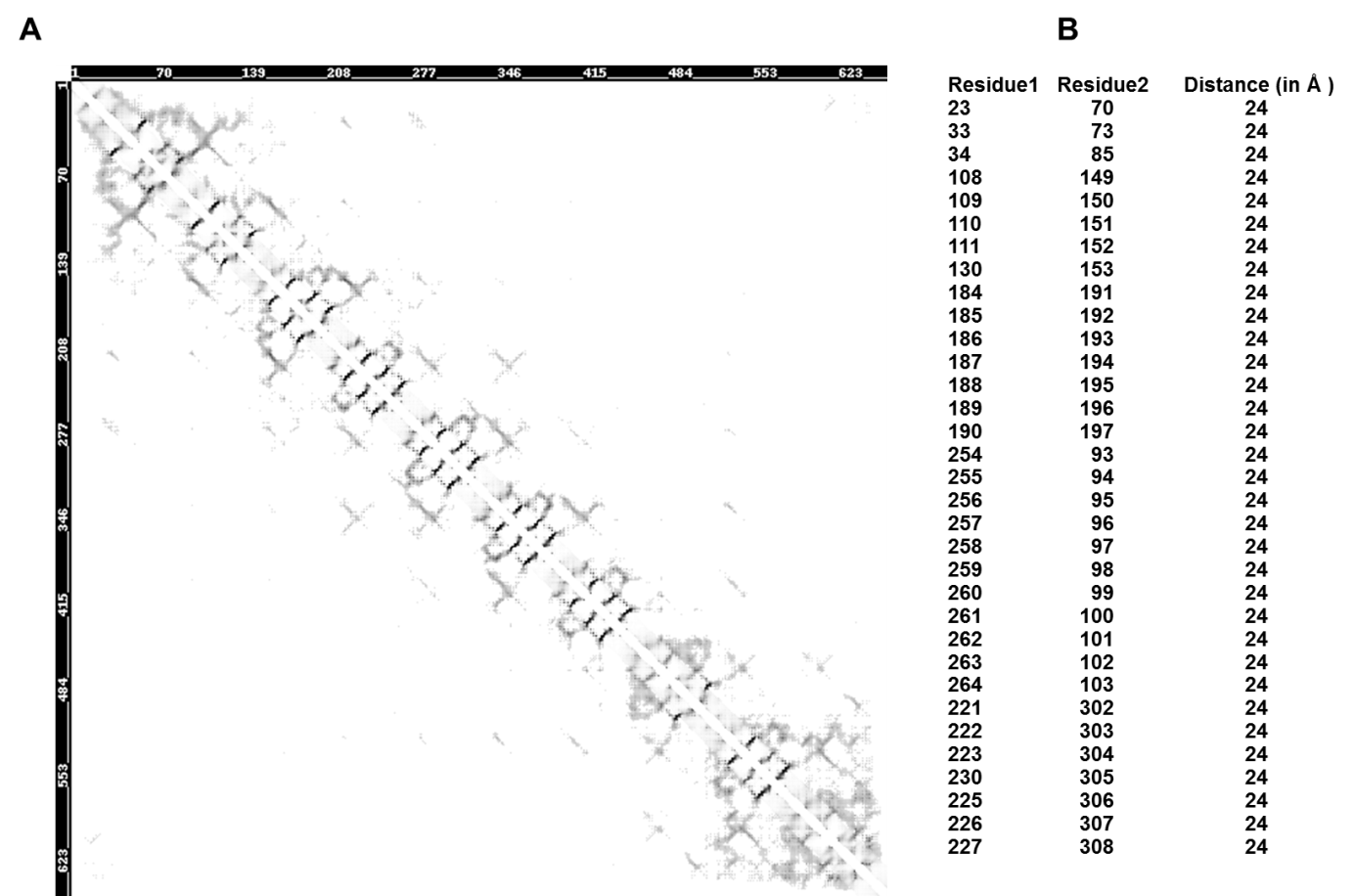

**Supplementary Figure 4** The different conformations of the FXIII-B monomer models (Models 1-5) assembled from their individual sushi domains by a combination of intra-subunit XL-MS derived constraints and contact residue prediction as well as assembled without any constraint (unbound) on the AIDA server. The Models 3-5 all colored blue did not result in successful/stable heterotetramer post docking downstream in this IH approach. All conformations are depicted in ribbon format. Model 1: assembled (on AIDA server) from XL-MS intra-subunit information + inter-residue contact prediction from Coinfold/RaptorX-Contact. Model 2: assembled from XL-MS intra-subunit information + inter-residue contact prediction from SVMSEQ. Model 3: assembled from XL-MS intra-subunit information + inter-residue contact prediction from PconsC3. Model 4: assembled from XL-MS intra-subunit information + inter-residue contact prediction from ICOS. Model 5: generated from XL-MS intra-subunit information + inter-residue contact prediction from CORNET. Unbound conformation model: Generated without any experimental constraints on the AIDA server. While both Model 1 and Model 2 generated stable heterotetramer on docking to FXIII-A2 crystal structure, Complex generated from Model 1 or Complex model 1 was the final chosen structure based on AFM data.

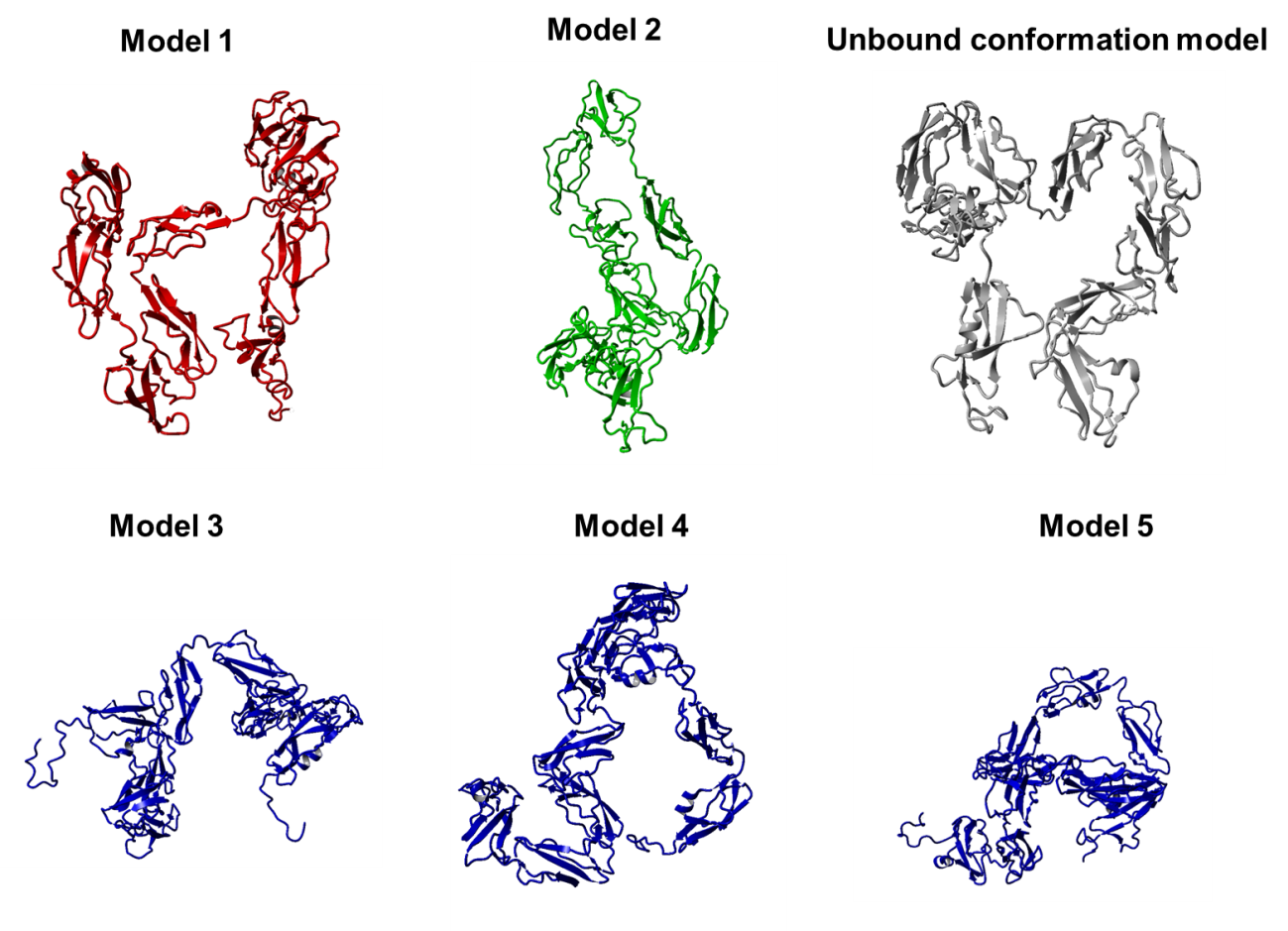

**Supplementary Figure 5a** Threaded modeling output files for FXIII-B subunit sushi domain 1 from ITASSER.

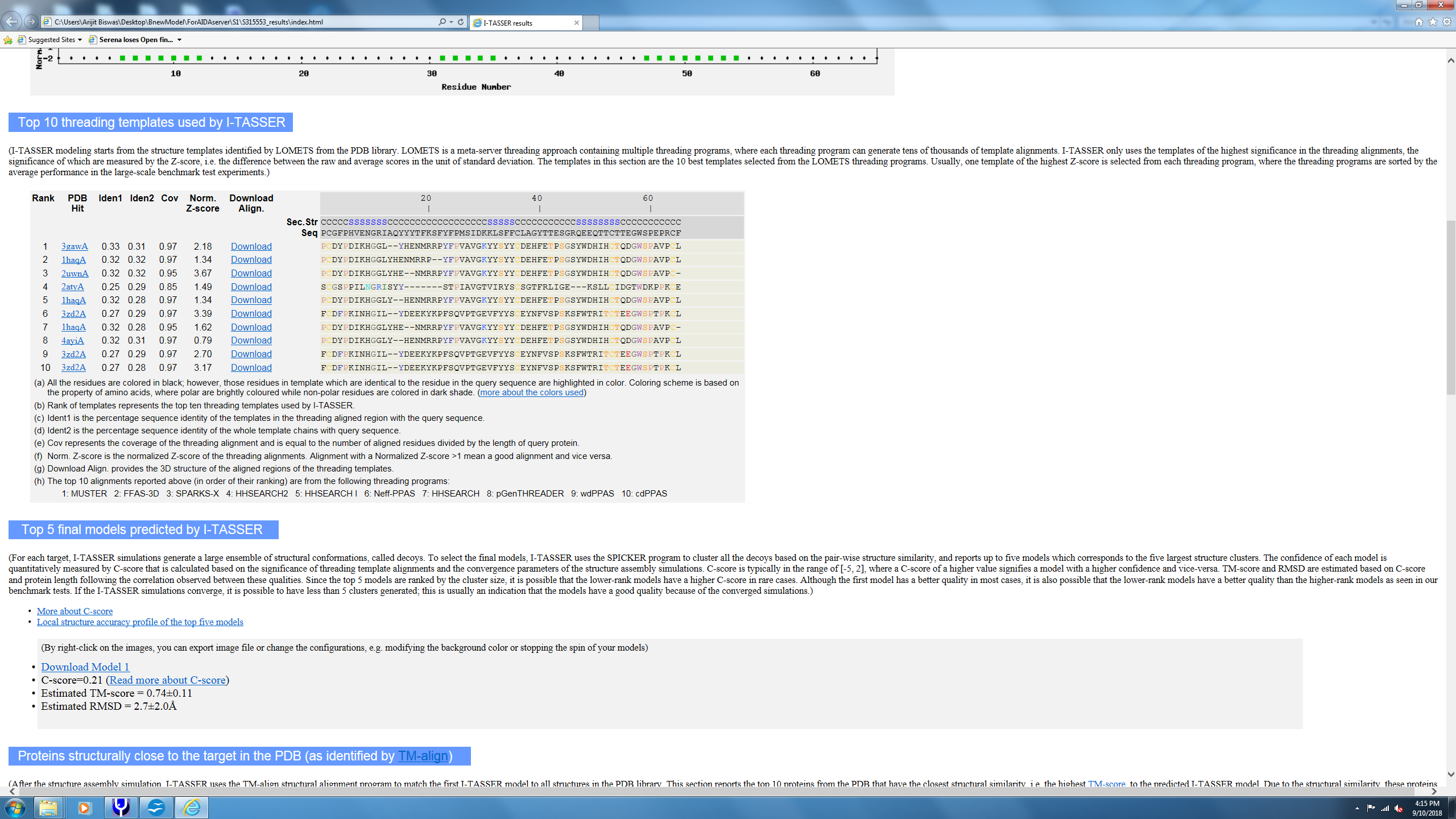

**Supplementary Figure 5b** Threaded modeling output files for FXIII-B subunit sushi domain 2 from ITASSER.

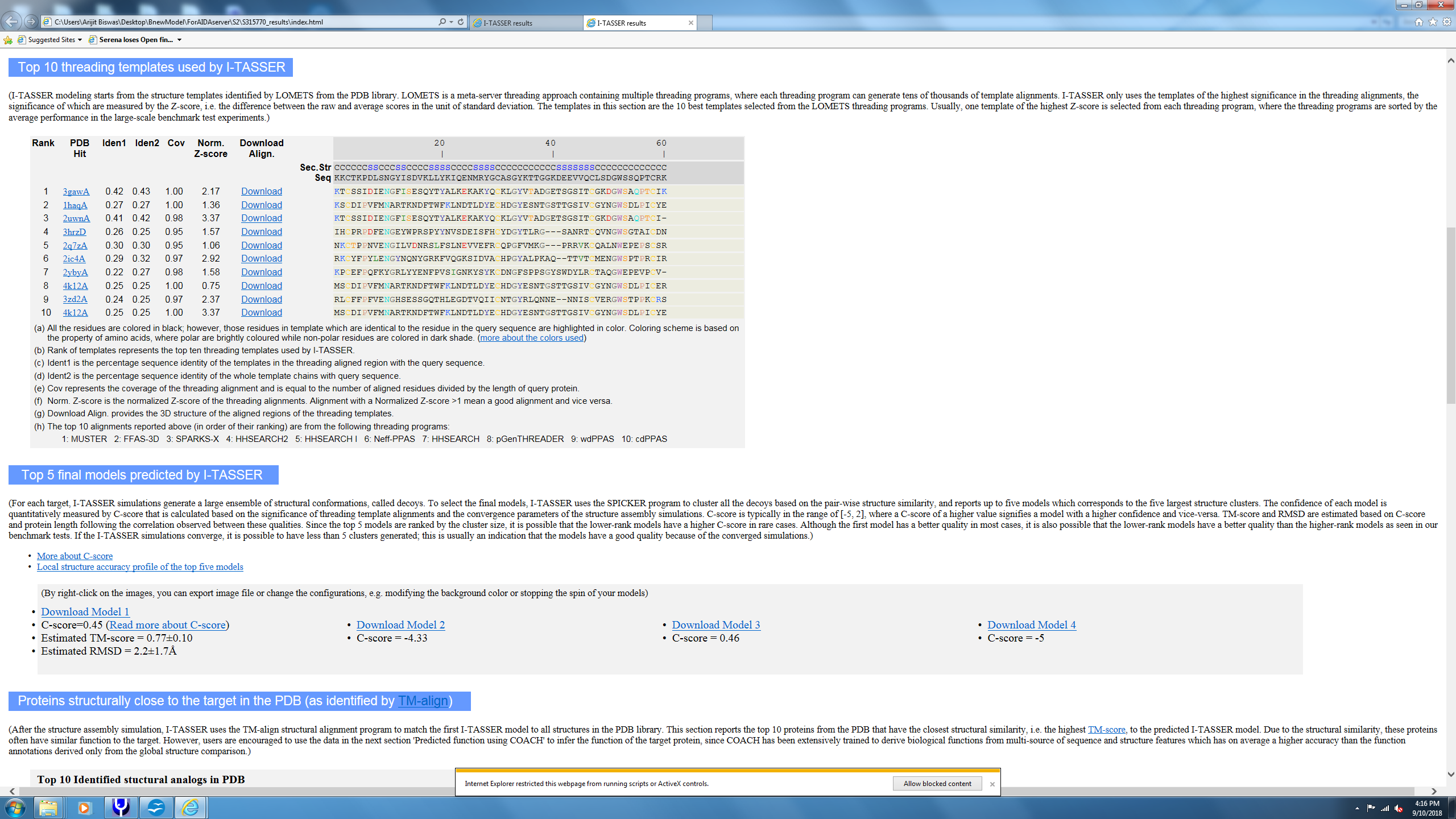

**Supplementary Figure 5c** Threaded modeling output files for FXIII-B subunit sushi domain 3 from ITASSER.

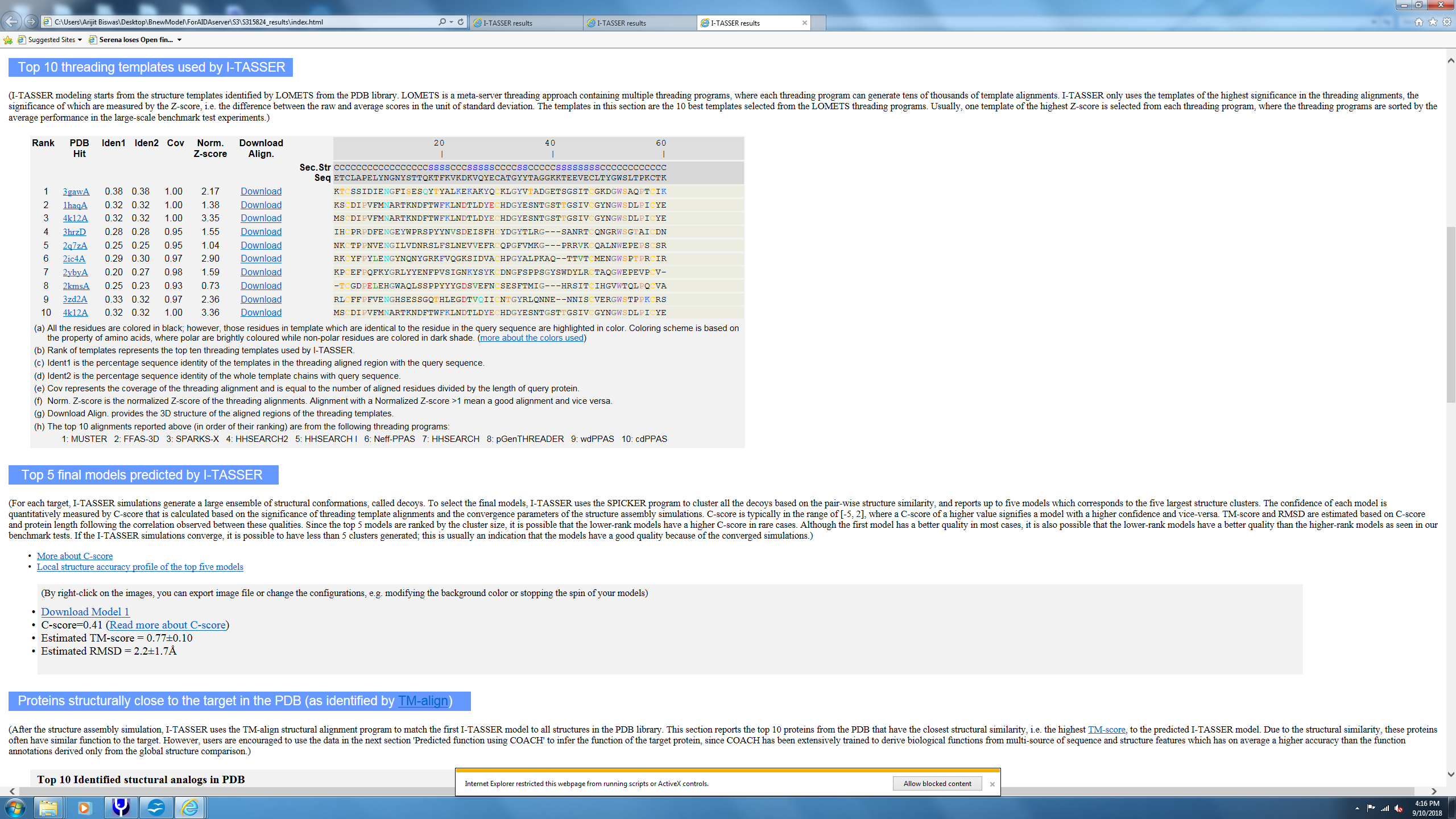

**Supplementary Figure 5d** Threaded modeling output files for FXIII-B subunit sushi domain 4 from ITASSER.

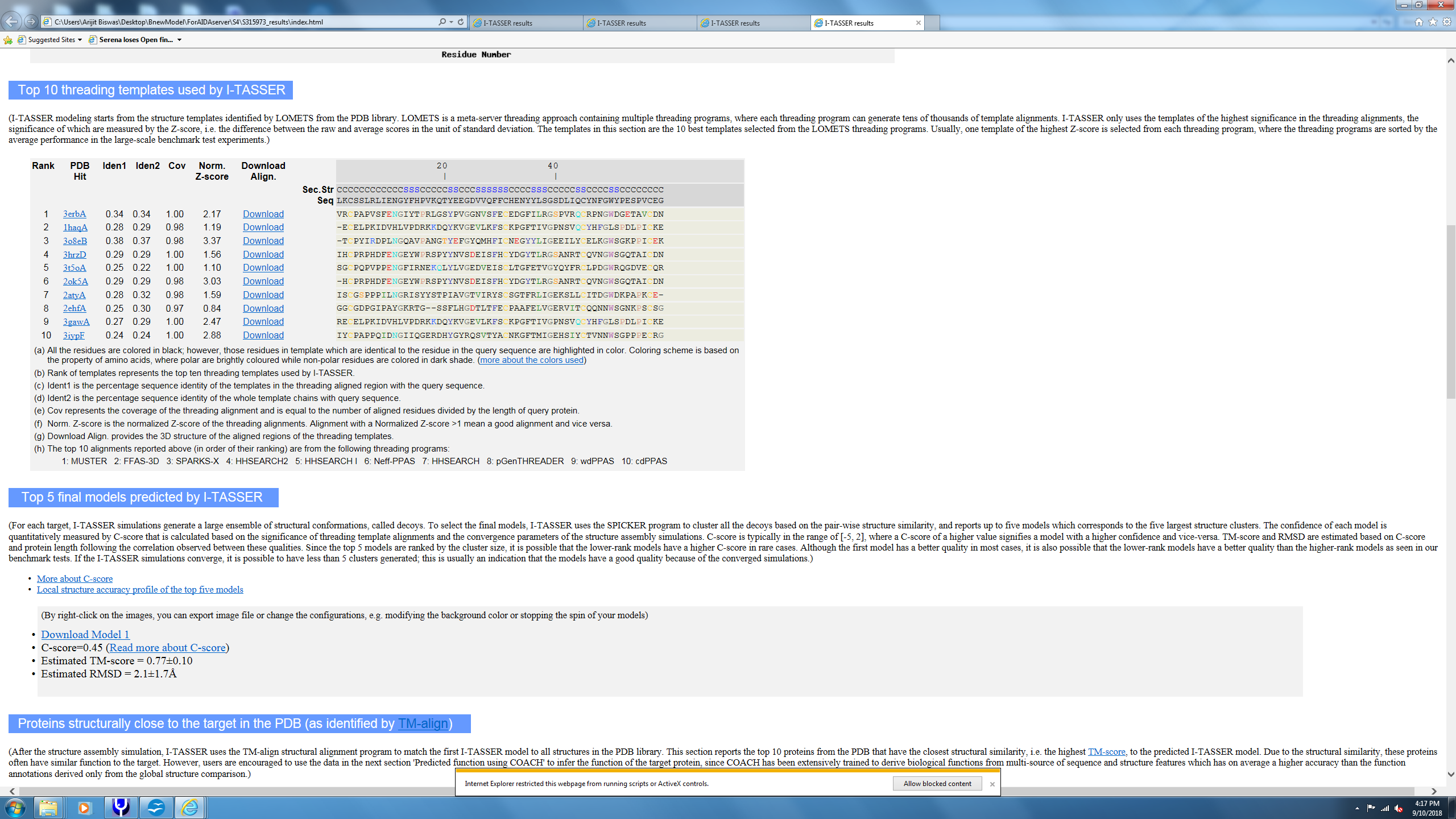

**Supplementary Figure 5e** Threaded modeling output files for FXIII-B subunit sushi domain 5 from ITASSER.

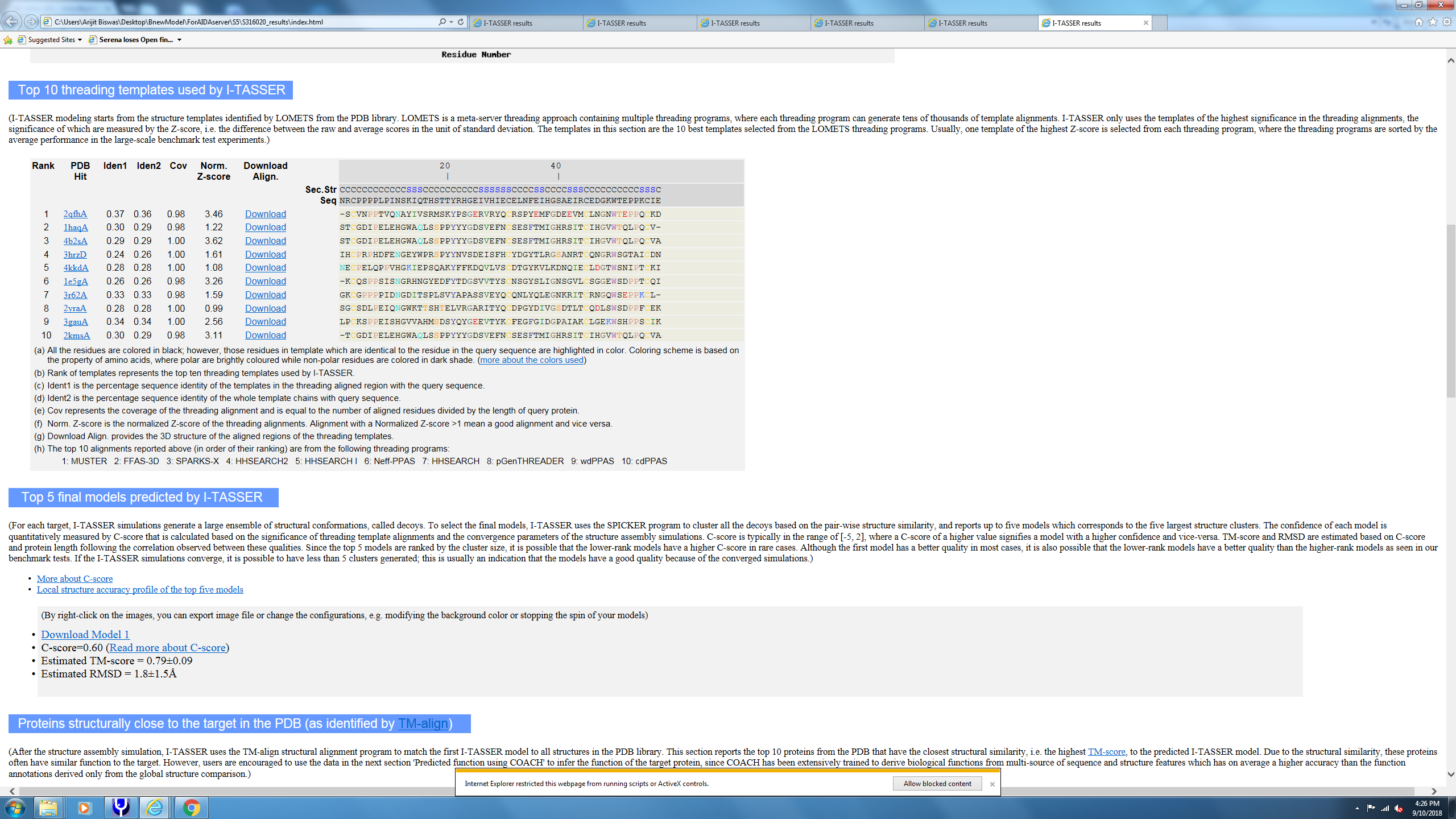

**Supplementary Figure 5f** Threaded modeling output files for FXIII-B subunit sushi domain 6 from ITASSER.

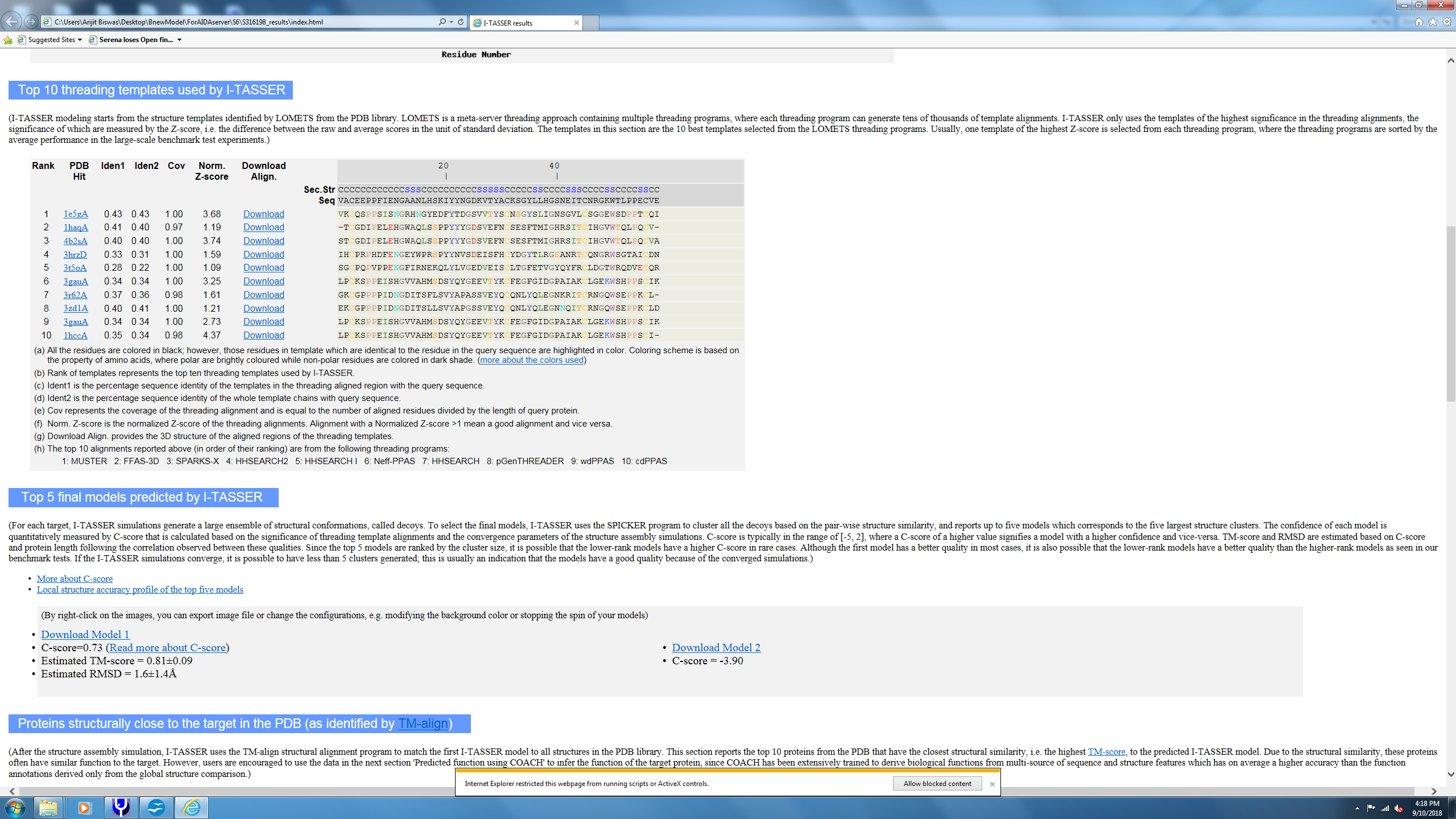

**Supplementary Figure 5g** Threaded modeling output files for FXIII-B subunit sushi domain 7 from ITASSER.

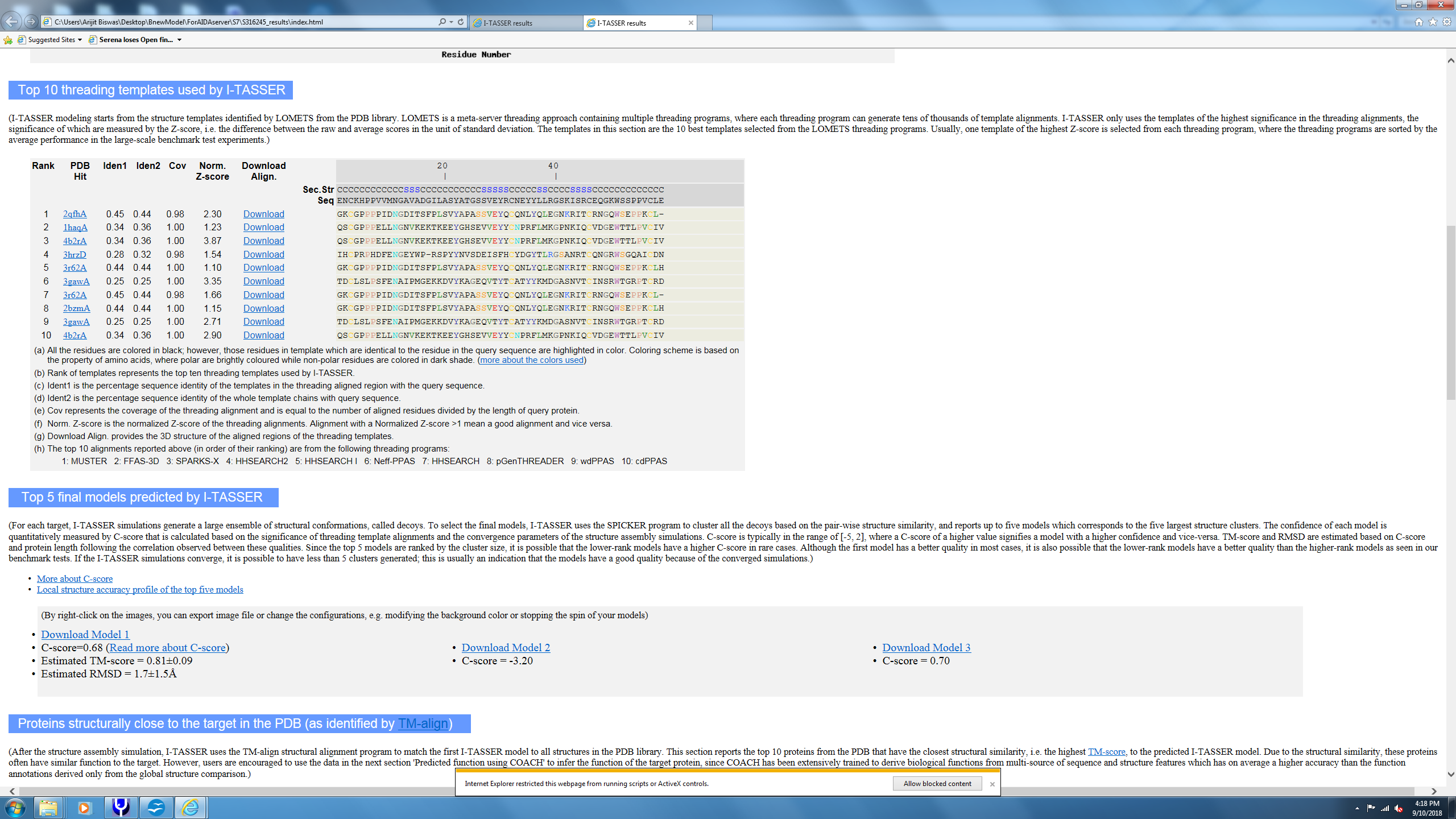

**Supplementary Figure 5h** Threaded modeling output files for FXIII-B subunit sushi domain 8 from ITASSER.

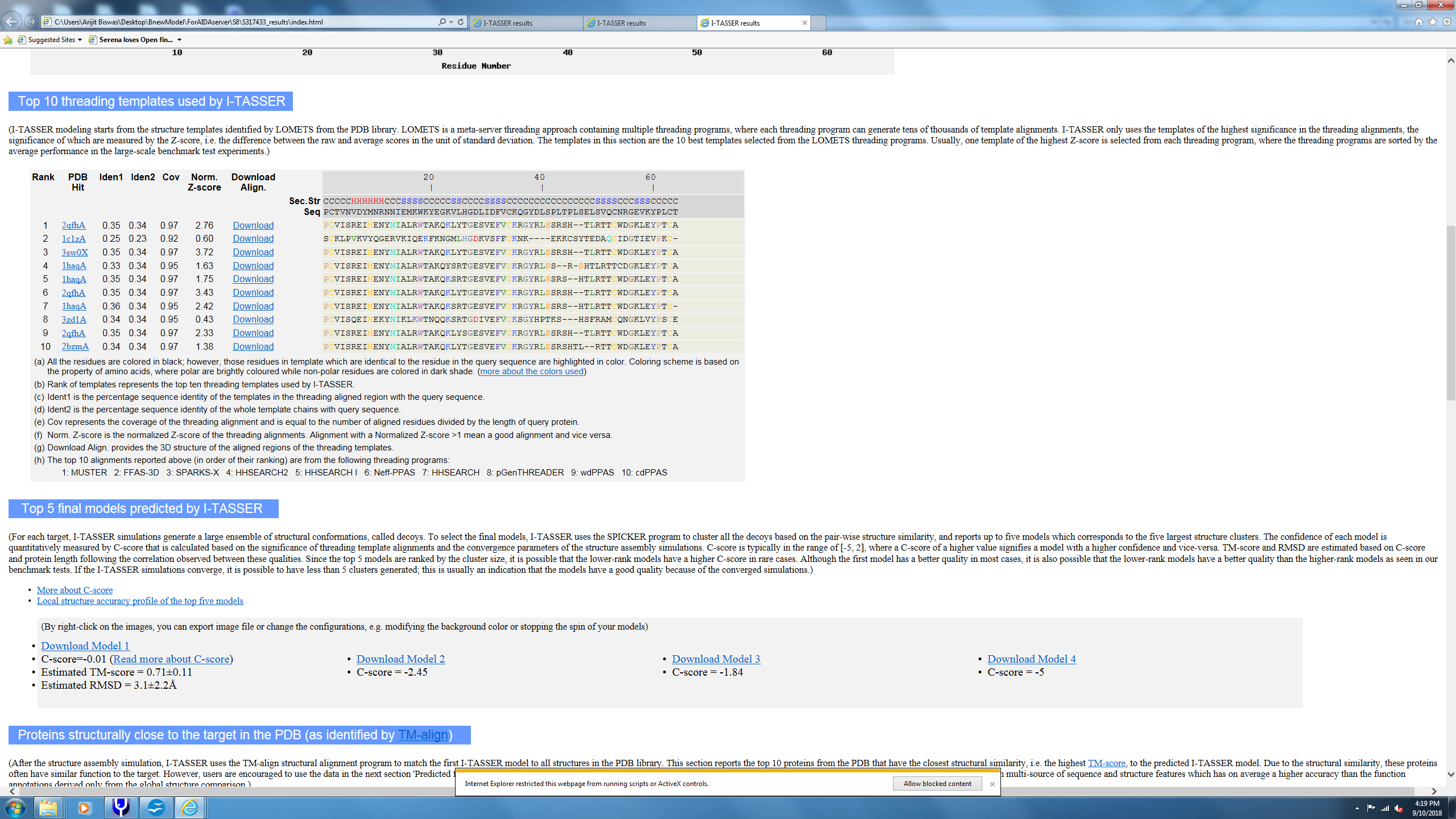

**Supplementary Figure 5i** Threaded modeling output files for FXIII-B subunit sushi domain 9 from ITASSER.

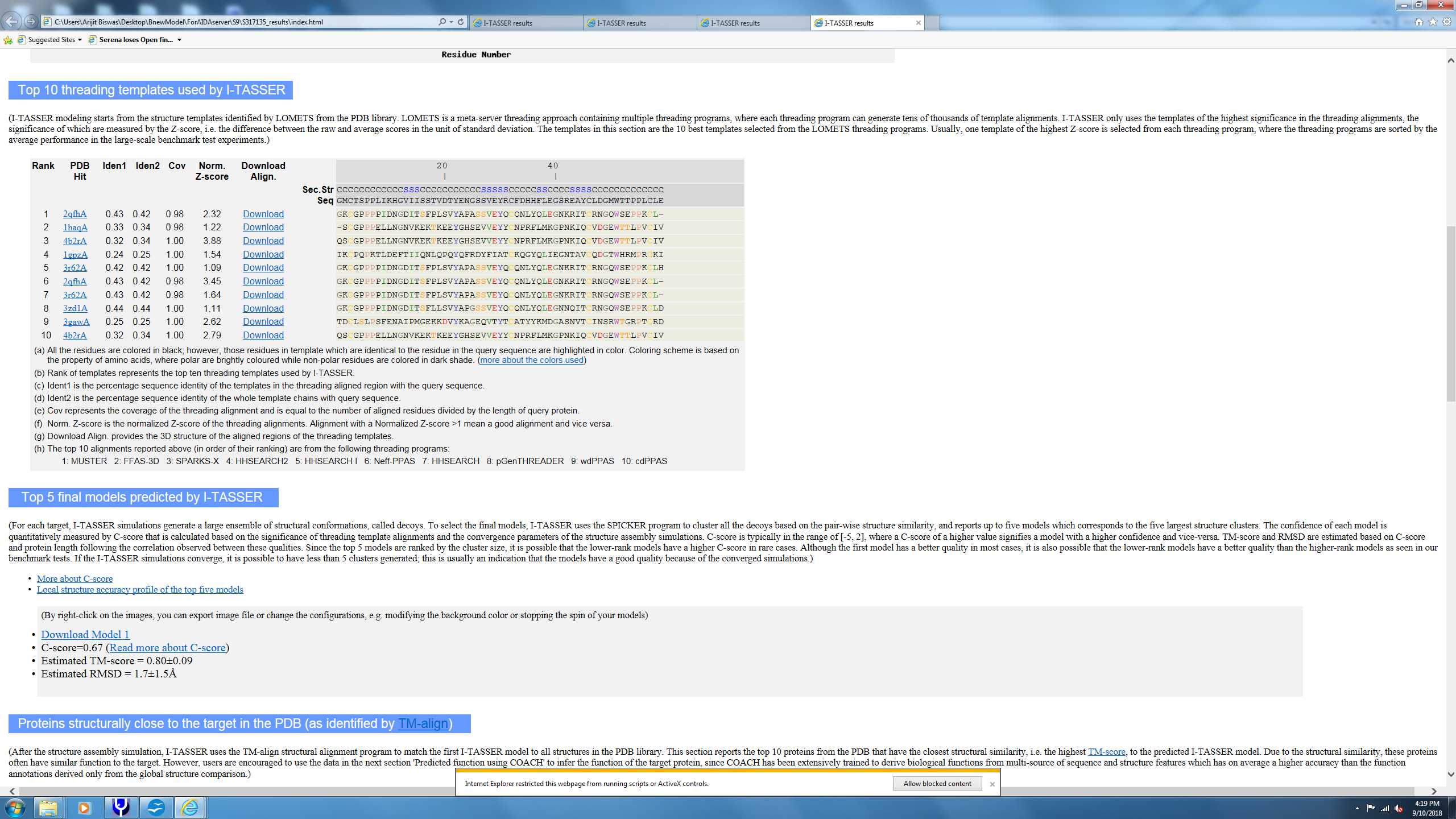

**Supplementary Figure 5j** Threaded modeling output files for FXIII-B subunit sushi domain 10 from ITASSER.

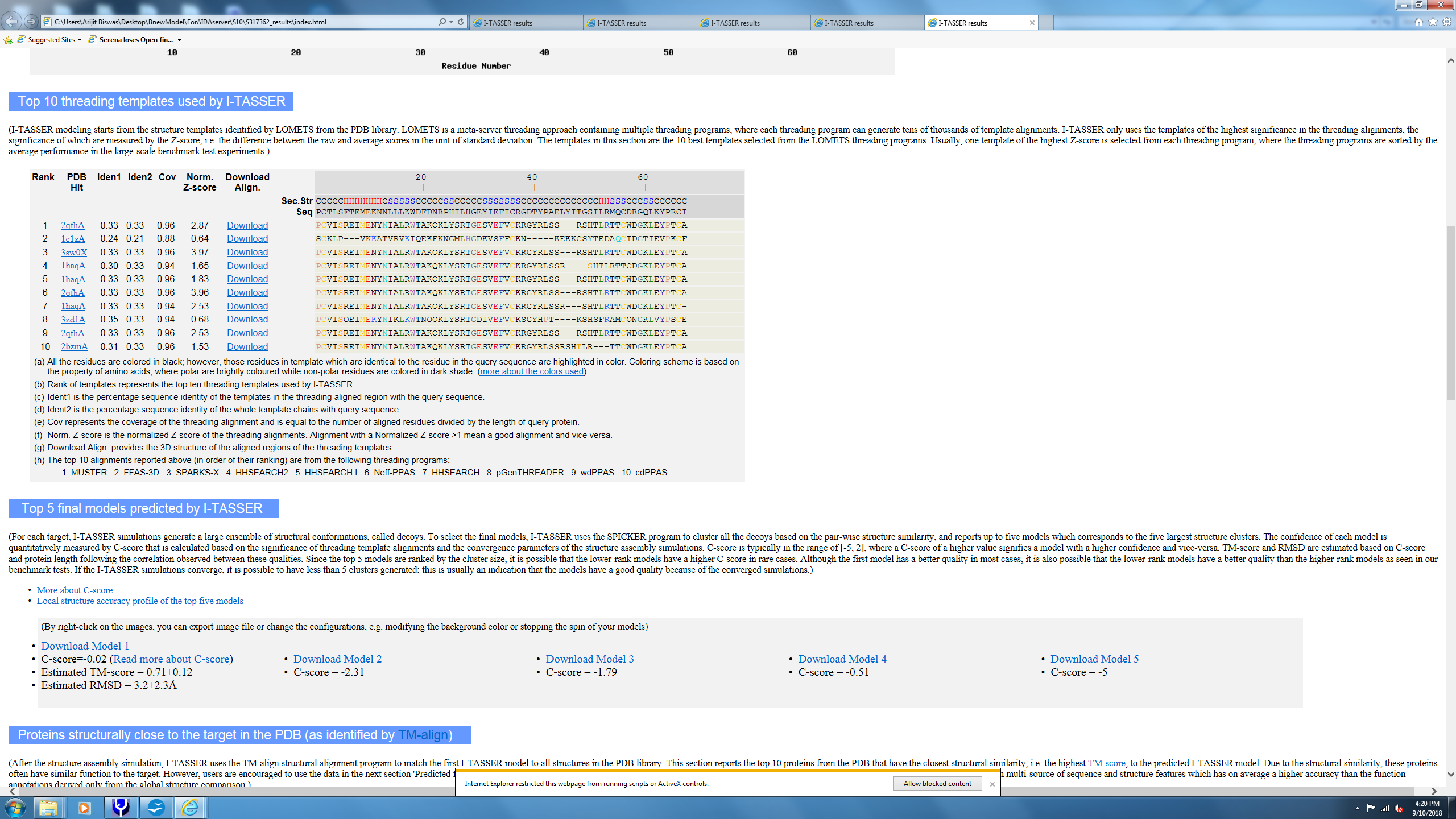

**Supplementary Figure 6** The HADDOCK scores for the flexible docking that generated model 1 of the FXIII complex. The scores are shown for all clusters resulting from the docking process. The model 1 was the best structure chosen from the top most cluster i.e. “cluster 1”.

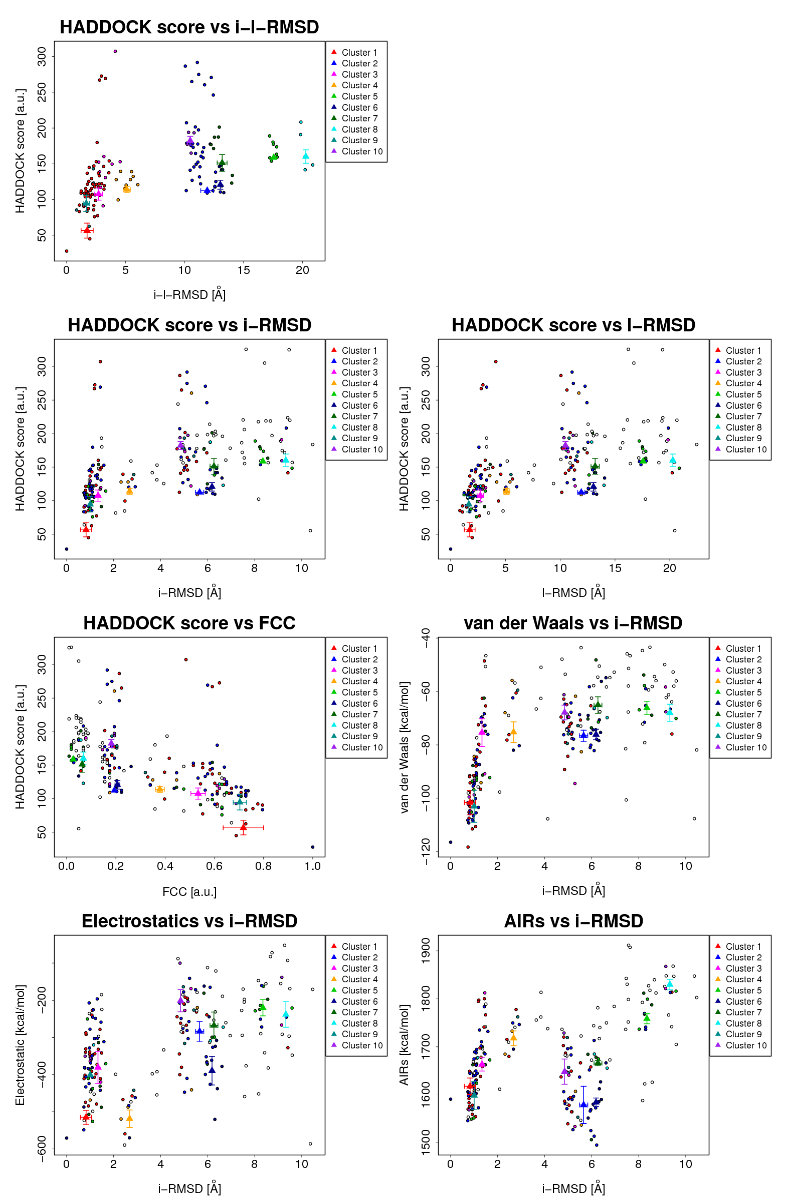

**Supplementary Figure 7** Close up view of topographic images of FXIII-A, FXIII-B subunits and of the FXIII complex obtained with AFM. The middle image shows one of the crops representing a FXIII complex with the bi-partite appearance and the flexible FXIII-B subunit peeking out from underneath the globular FXIII-A subunit.

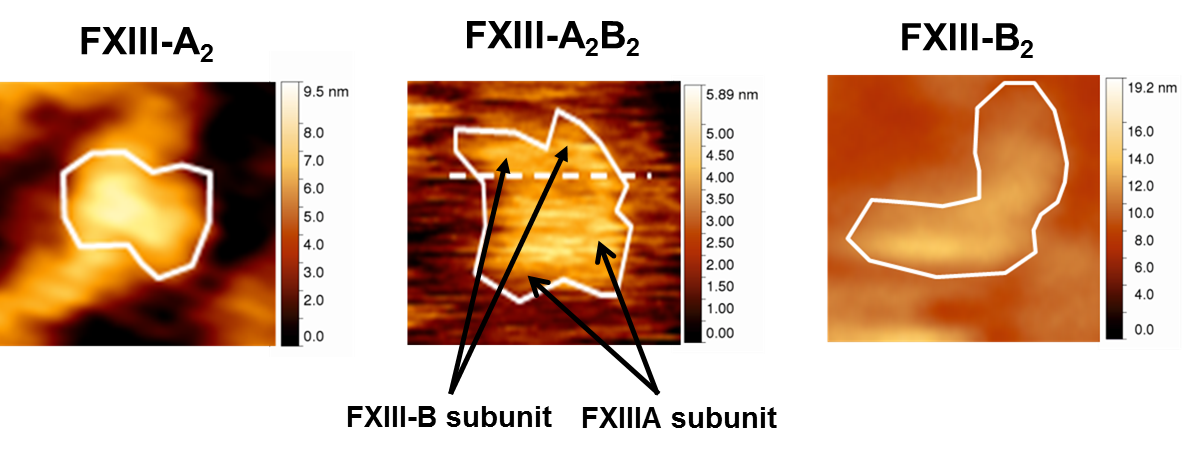

**Supplementary Figure 8** The structures of the two stable all atom models of the FXIII-A_2_B_2_ successfully generated from the flexible docking of monomeric FXIII-B subunit models. These models represent the topmost model (based on HADDOCK scores) of the best docking cluster from each of the two successful docking exercises.

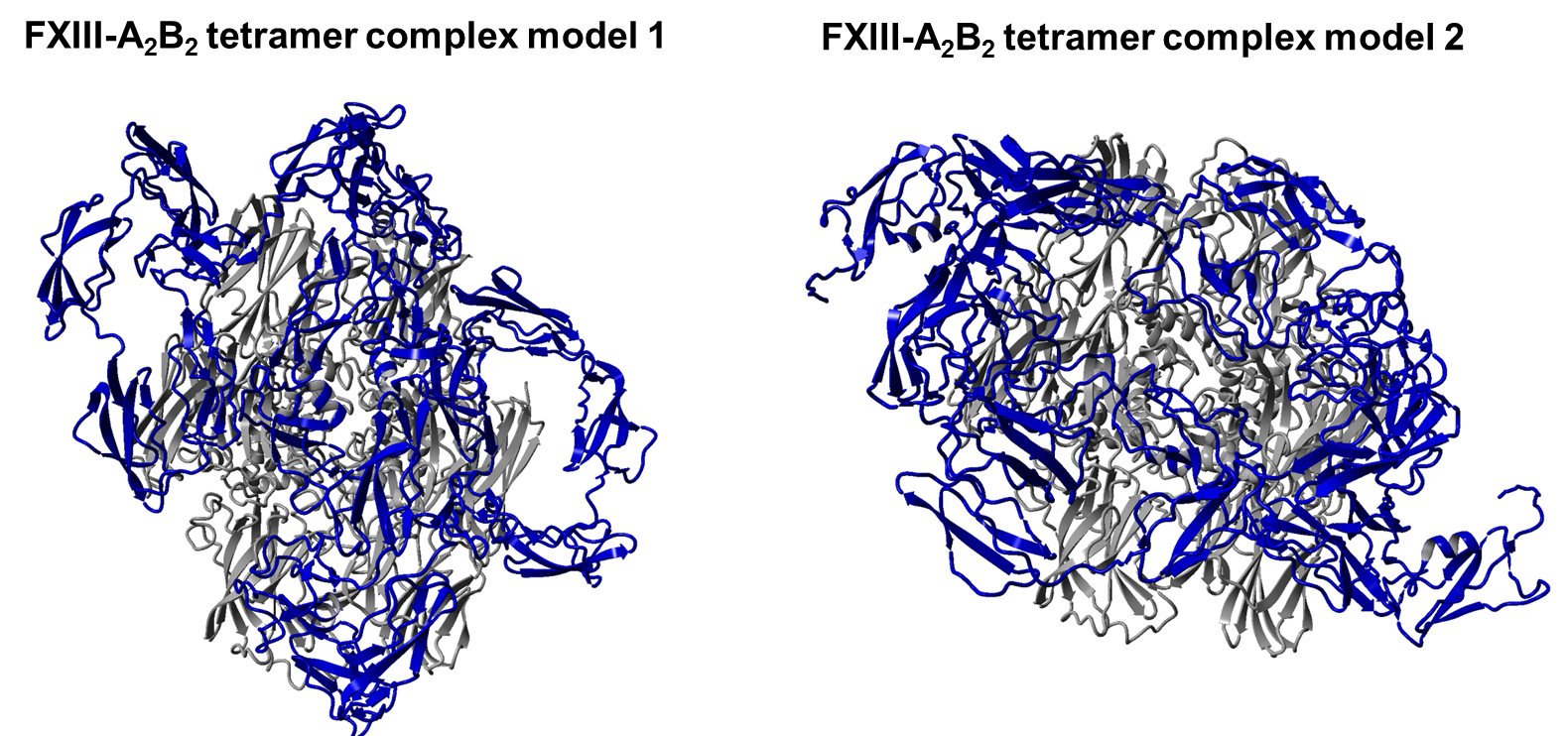

**Supplementary Figure 9** The stereochemical validation output for model 1 of the heterotetrameric FXIII-A_2_B_2_ complex structure obtained from the SAVES v5.0 server.

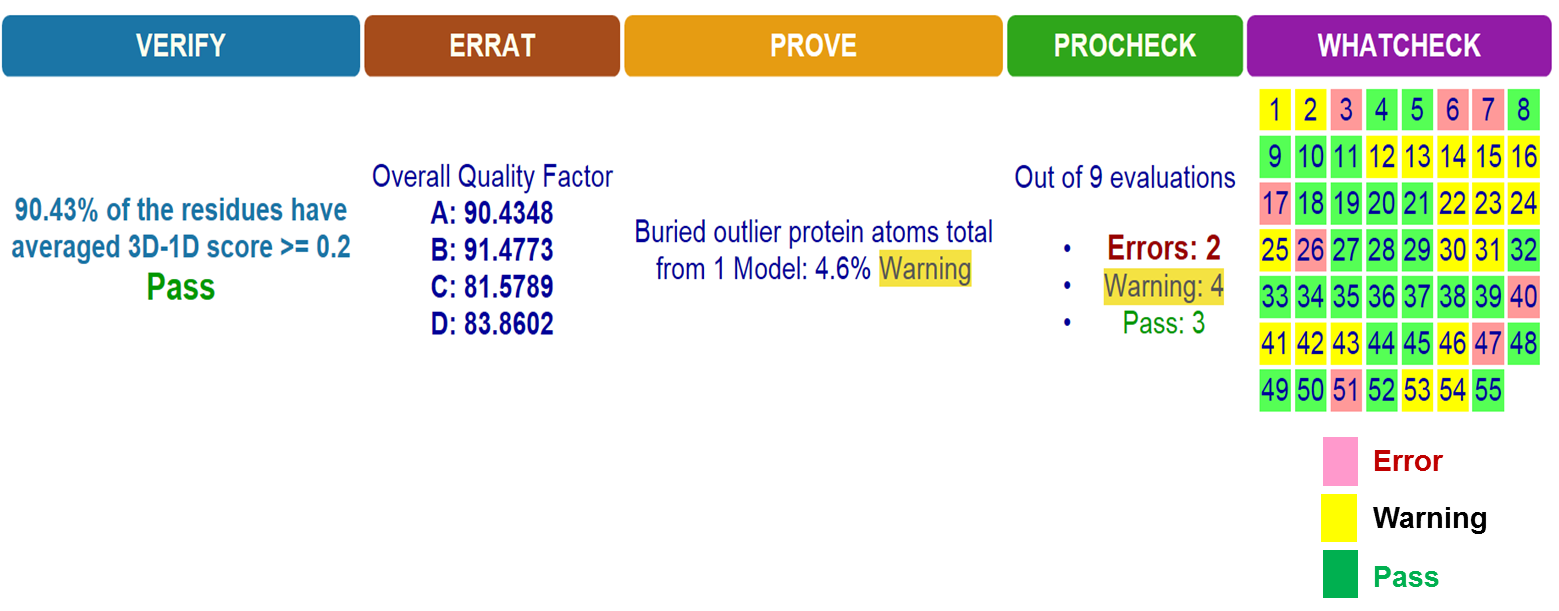

**Supplementary Figure 10** Two models for the assembly (association) and disassembly (during activation of the FXIII-A and FXIII-B subunits.

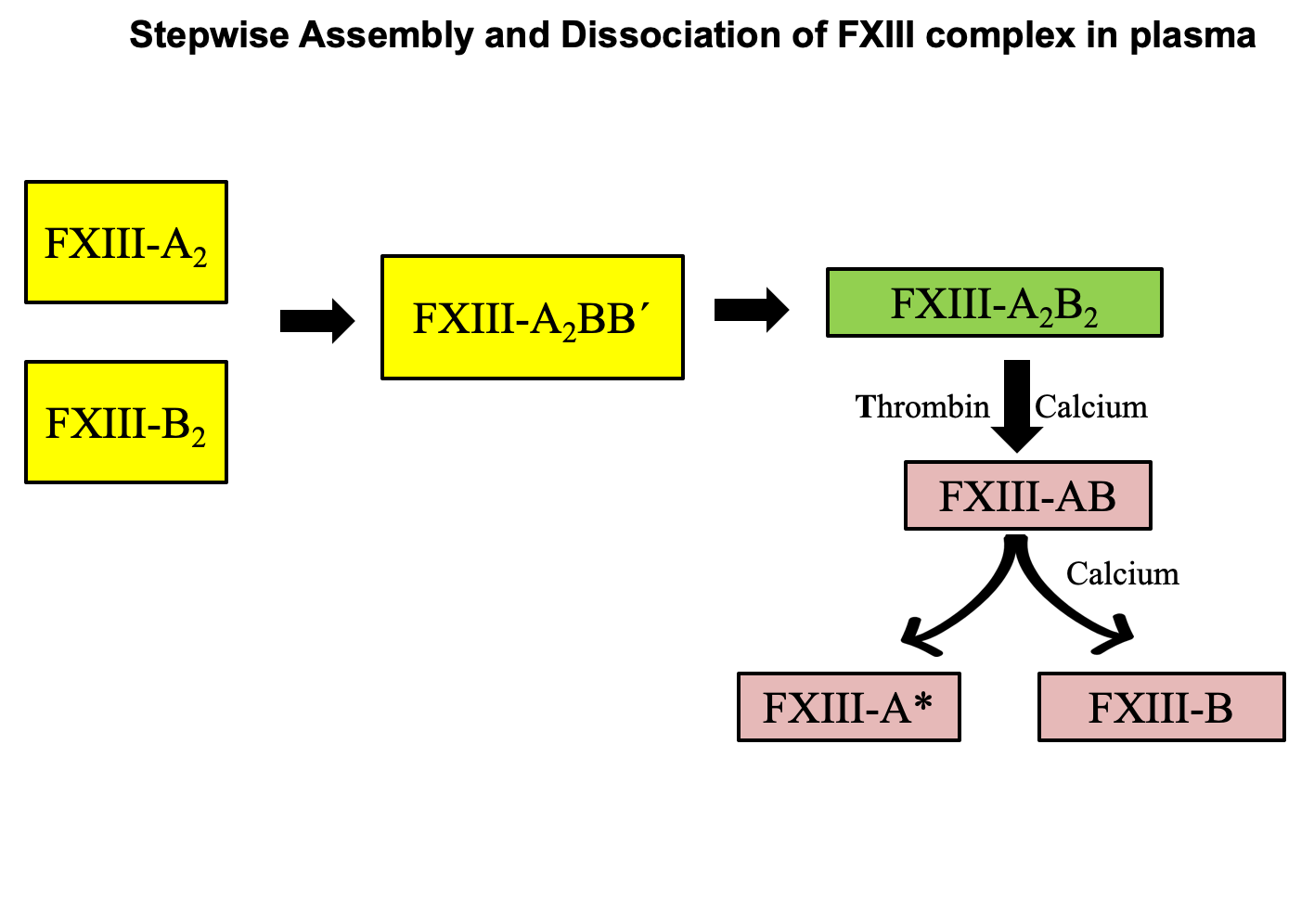

**Supplementary Figure 11a** shows a structural alignment between the zymogenic and non-proteolytically activated (FXIII-Aa) monomeric structures of FXIII-A subunit in cyan and green colored ribbon format respectively. The disordered hinge region around which the barrel domains pivot during the activation of FXIII is shown in blue. **Supplementary Figure 11b** shows the molecular surface of the heterotetramer complex model 1 with the molecular surface of FXIII-B subunit colored cyan and that of FXIII-A subunit colored grey. The disordered hinge region in the FXIII-A subunit shown in **Supplementary Fig 11a** is no longer disordered owing to its interaction with FXIII-B subunit in one monomer as shown by the inset structure. However, on the other monomer the region is still disordered owing to unequal pairing and lack of interaction with the FXIII-B subunit allowing for the entry of bulk solvent.

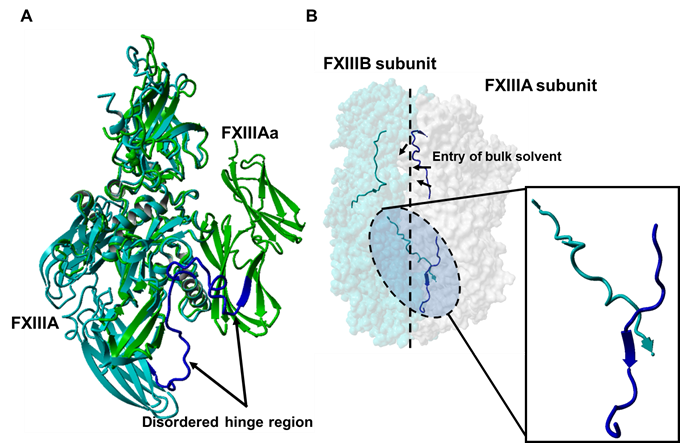

**Supplementary Figure 12** The steered simulated dissociation of FXIII-B_2_ subunit from the FXIII-A_2_B_2_ complex model

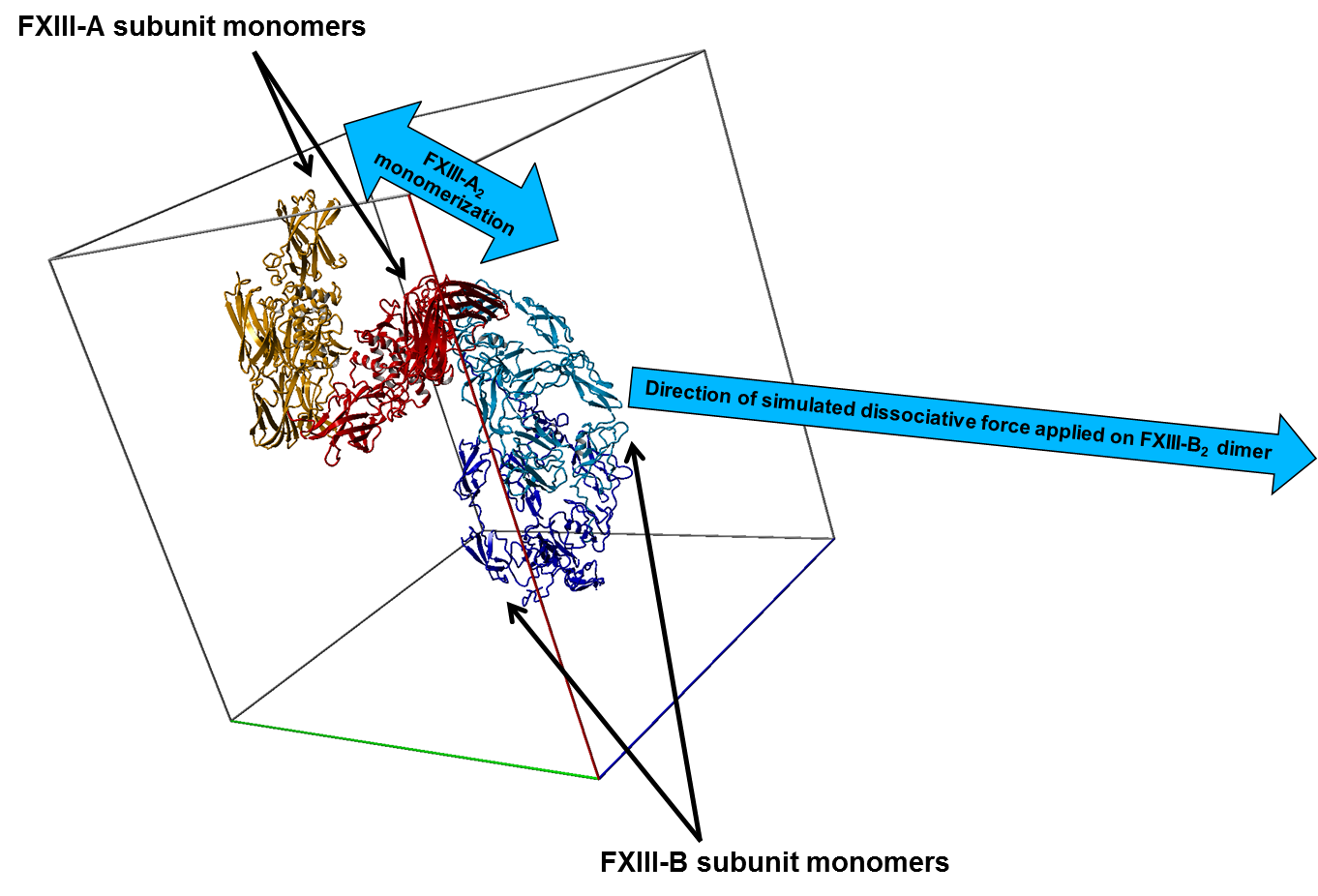

**Supplementary video legends**

**Supplementary Video 1.** This video shows the thermal motion of the heterotetramer complex model 1 for a short period of the MD simulation. The FXIII-A subunit chains are depicted in red and orange colored ribbon formats. The FXIII-B subunit chains are depicted in blue and cyan colored ribbon formats.

**Supplementary Video 2.** This video shows the simulated dissociation of the FXIII-B subunit dimer from the FXIII-A subunit dimer within the heterotetramer complex model 1. The FXIII subunit chains are depicted in similar format to Supplementary Video 1.

**Supplementary video 3.** This video shows a close up view of thermal motion of disordered flexible hinge region from the FXIII-A subunit (residues 500-520) as observed in the MD simulation of the FXIII-A subunit dimer crystal structure. The structure of this region is depicted in grey colored ribbon format.

**Supplementary video 4.** This video shows a close up view of thermal motion of the interface between C-terminal FXIII-B subunit and the otherwise disordered flexible hinge region from the FXIII-A subunit (residues 500-520) as observed in the MD simulation of the heterotetramer complex model 1. The structure of the FXIII-B subunit is depicted in yellow colored ribbon format while the FXIII-A subunit is depicted in grey colored ribbon format

1. *^*^* ***Corresponding author:***

   PD Dr. Arijit Biswas,

   Institute of Experimental Hematology and Transfusion Medicine, Sigmund-Freud Street 25, University Hospital of Bonn, 53127 Bonn, Germany. Phone: +49-22828719428; Fax: +49-22828716087; E-Mail:
